# Spiking networks that efficiently process dynamic sensory features explain receptor information mixing in somatosensory cortex

**DOI:** 10.1101/2024.06.07.597979

**Authors:** Veronika Koren, Alan J. Emanuel, Stefano Panzeri

**Affiliations:** Institute of Neural Information Processing, Center for Molecular Neurobiology (ZMNH), University Medical Center Hamburg-Eppendorf (UKE), 20251 Hamburg, Germany; Department of Cell Biology, Emory University School of Medicine, Atlanta, GA, 30322, USA; Istituto Italiano di Tecnologia, Genova, Italy

**Keywords:** efficient coding, spiking neural networks, recurrent neural networks, somatosensory system

## Abstract

How do biological neural systems efficiently encode, transform and propagate information between the sensory periphery and the sensory cortex about sensory features evolving at different time scales? Are these computations efficient in normative information processing terms? While previous work has suggested that biologically plausible models of of such neural information processing may be implemented efficiently within a single processing layer, how such computations extend across several processing layers is less clear. Here, we model propagation of multiple time-varying sensory features across a sensory pathway, by extending the theory of efficient coding with spikes to efficient encoding, transformation and transmission of sensory signals. These computations are optimally realized by a multilayer spiking network with feedforward networks of spiking neurons (receptor layer) and recurrent excitatory-inhibitory networks of generalized leaky integrate-and-fire neurons (recurrent layers). Our model efficiently realizes a broad class of feature transformations, including positive and negative interaction across features, through specific and biologically plausible structures of feedforward connectivity. We find that mixing of sensory features in the activity of single neurons is beneficial because it lowers the metabolic cost at the network level. We apply the model to the somatosensory pathway by constraining it with parameters measured empirically and include in its last node, analogous to the primary somatosensory cortex (S1), two types of inhibitory neurons: parvalbumin-positive neurons realizing lateral inhibition, and somatostatin-positive neurons realizing winner-take-all inhibition. By implementing a negative interaction across stimulus features, this model captures several intriguing empirical observations from the somatosensory system of the mouse, including a decrease of sustained responses from subcortical networks to S1, a non-linear effect of the knock-out of receptor neuron types on the activity in S1, and amplification of weak signals from sensory neurons across the pathway.

## 1 Introduction

Fundamental computations of neural information processing include the steps of encoding, transformation and transmission of information needed for the generation of behavior. An influential hypothesis used to understand the principles behind these fundamental computations is the efficient coding hypothesis ^1^, which posits that neural systems maximize information about external stimuli while at the same time controlling the metabolic cost of spiking activity ^2^. This hypothesis has been first used to study principles of computations in artificial neural networks lacking biological realism such as spiking activity ^3^. Recent advances have extended the normative theory of efficient coding to implementations with models of spiking neural networks ^4,5^ including biologically plausible spiking networks with excitatory (E) and inhibitory (I) neurons ^6,7^. A limitation of these and other previous studies on efficient coding ^8,9,10,11,12^ (but see ^13^) is that it has focused on information processing within a single layer of neurons. In biological brains, sensory information travels across a number of brain areas that can be conceptualized as a succession of neural processing layers. This implies that any principled theory of neural information processing must include information transmission ^14,15,16^ and potential transformation of sensory signals across processing layers. An important question is, how does the brain efficiently encode, transform and transmit correlated sensory signals on different time scales between sensory periphery and sensory cortex with biologically plausible building blocks. Another question is what are the computations relevant to biological sensory pathways.

To address these questions, we extended the theory of efficient coding with spikes and built a multilayer spiking network that optimizes encoding, transformation and transmission of sensory signals, with a focus on the ascending somatosensory system of the mouse. Our approach combines analytical, optimization-based approach with the knowledge of constraints and properties of biological neural networks, such as Dale’s principle, distributed neural code ^17,18^, convergence of receptor neurons ^19^ and the architecture of the FF connectivity ^20^ that are suggested by empirical studies. Efficient encoding stems from minimizing layer and neuron type specific encoding error between targeted and actual network representations with limited numbers of spikes. Efficient transformation and transmission is based on the assumption that the function of sensory information from the previous layer defines a target signal of the next layer. This assumption leads to specific structure of the feedforward connectivity between the two connecting processing layers recovered analytically.

In this paper, we first develop an analytical theory of the efficient multilayer spiking network that we built from normative principles of efficient coding and constraints from neurobiology. We then test the theory by simulating a simplified but general model of sensory processing with optimally efficient parameters and demonstrate how its emerging properties benefit efficient coding. We finally present a more complex network with two types of inhibition in its last layer and with parameters constrained with biophysics. We show that this model captures non-linear mixing of fast and slow features of tactile receptors in the somatosensory cortex as it implements a negative interaction across these features.

## 2 Results

### 2.1 Building an efficient sensory pathway from efficient coding theory and biological constraints

Somatosensory information enters the nervous system through receptors in the skin. We denote the entry level of processing as L0 and we assume that we have *m* = 1, …, *M* types of sensory receptor neurons, with each receptor specialized to respond to one feature of the stimulus ^21^. The way a receptor neuron responds to a stimulus depends on its molecular and structural properties differs widely across receptor neuron types. We capture this diversity by defining features of the stimulus 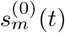 that receptors of type *m* are sensitive to. We call the computation that sensory neurons of type *m* perform on their stimulus feature the target representation and denote it with 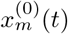. We assume that this computation is a leaky integration of the stimulus feature 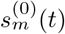. The feature and the target representation are identical across receptor neurons of the same type and the superscript ^(0)^ means that these neurons belong to L0.

We further assume that the activity of each sensory neuron *i* minimizes an efficiency objective that evaluates the encoding error and the metabolic cost of neuron’s activity:

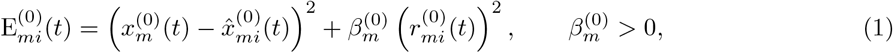

with 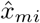 a dynamical readout and *r*_*mi*_ the low-pass filtered spike train of the neuron and 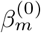 the weighting between the encoding error and the metabolic cost. By assuming that the neuron emits a spike only if this momentarily decreases its loss function, we analytically derive a generalized leaky IAF neuron with feedforward and adaptation currents as an exact solution to the optimization problem in Eq. 1 (see Appendix A.1.1). These derivations, as well as all other theoretical proofs, involve linear algebra and differential equations. We find that the feedforward current to the neuron is proportional to neuron’s preferred sensory feature 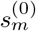.

We apply our theory to the mammalian somatosensory system and to processing of stimuli that evoke the perception of fine touch. Such stimuli evoke responses from two types of skin receptors, slowly and rapidly adapting low threshold mechanoreceptors (SA-LTMRs and RA-LTMRs, ^22,23,24^). We design stimulus features to each receptor neuron type to capture the impulse response profiles of these neurons observed empirically in the somatosensory system of the mouse ^25^. Our analytically derived model of L0 has a handful of free parameters that we constrained with computer simulations, such that the simulated model minimized a weighted sum of the average encoding error and metabolic cost (see Appendix A.3) in response to a naturalistic stimulus (Ornstein-Uhlenbeck process, see Appendix A.5). Optimal parameters of receptor neurons were within biophysical ranges expected for somatosensory receptors (Fig. S1 A-B). The output of SA-LTMRs and RA-LTMRs accurately tracked the time-dependent fluctuations of the target representation (but not its mean and variance), with the coefficient of determination on z-scored signals of 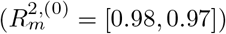 for SAand RA-LTMRs, respectively. Such optimal networks of sensory receptors activated with biologically plausible activity profiles and with high temporal sensitivity (Fig. 1C, Fig. S1 C-D).

**Figure 1.**
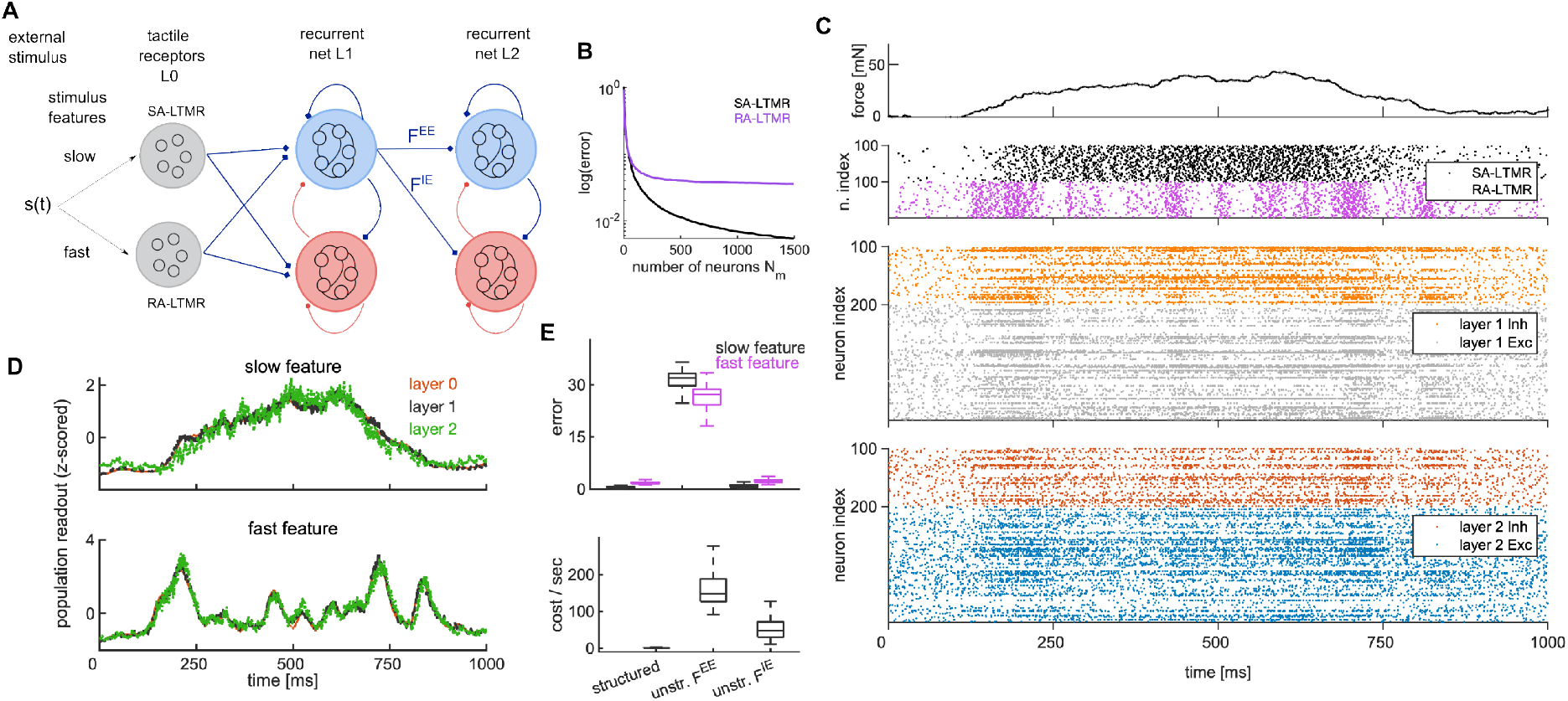
Multilayer efficient spiking network accurately propagates a slow and fast feature of the stimulus across layers. **A** Schematic of the model. From the external stimulus *s*(*t*), receptor neurons of type SA-LTMR and RA-LTMR extract a slow and a fast stimulus feature that are transmitted to E (blue) and I (red) neurons in recurrently connected layers L1-2. **B** Log of the encoding error as a function of the number of receptor neurons. **C** Spiking activity in an example simulated trial. Top: external stimulus (force on the skin). 2nd from top: spikes of receptor neurons SA-LTMR (black) and RA-LTMR (purple). 3rd from top: spiking activity of E (gray) and I (orange) neurons in L1. Bottom: spikes of E (blue) and I (red) neurons in L2. Parameters are identical for L1-2. **D** Population readout of the slow (top) and fast feature (bottom) across layers in one simulation trial. **E** Error (top) and cost per second (bottom) in L1-2 with and without convergence of sensory signals. Significance is assessed with two-tailed t-test and Bonferroni correction. See SM Table S1–S2 for parameters.

We next define the transmission of information between L0 and the next processing layer, L1. Empirical studies of early somatosensory processing have emphasized convergence of sensory neurons on their postsynaptic targets ^19,26^, where multiple receptor neurons of both types have synapses on the same neurons in the downstream area. To achieve such convergence, we define the feedforward synaptic current from L0 to L1 as a linear sum of spiking activity across receptor neuron types as well as across neurons of the same type:

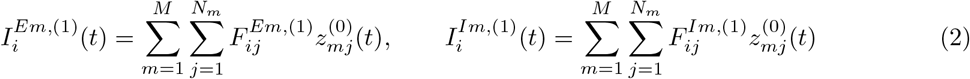

With 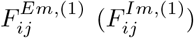 the synaptic weight between the presynaptic (receptor) neuron *j* (of type *m*) and the excitatory (E) and inhibitory (I) postsynaptic neuron *i* in L1. The variable 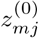 is the low-pass filtered spike train of a receptor neuron. With FF current between L0 and L1 as in Eq. 2, we achieve convergence of information about fast and slow sensory features on single neurons in L1. We find that within-type convergence of receptors improves information transmission, as the error between the target signal and the across-neuron averaged readout decreases by increasing the number of converging receptor neurons in both receptor neuron types (Fig. 1B).

In spite of convergence of sensory information onto single neurons, it may be desirable for the L1 network to disambiguate between neural signals encoding fast and slow feature to allow precise computations on stimulus features downstream. To achieve this, we endow the target signal in L1, **x**^(1)^(*t*), with several dimensions. In particular, the dimensionality of **x**^(1)^(*t*) as well as of all target signals beyond L1 is given by the total number of receptor neuron types *M*. In our model of the somatosensory system, the first and second dimension capture the information about the slow and fast feature, respectively, while the remaining dimensions are unused and provide an equivalent to a representational noise (Fig. S4A). Our model of the FF connectivity between L0 and L1 gives rise to efficient transmission of the information about sensory features across these two layers and allows the co-existence of convergence of sensory features on single neurons in L1, as well as the possibility to decode both features from the activity of the L1 network. We assume that no transformation on sensory features takes place in L1.

To account for transformations in the representation of the sensory information, we assume that the target signal in a generic layer L(n), **x**^(*n*)^(*t*) with *n* ≥ 2, depends on a linear transformation of the population readout from the previous (presynaptic) layer L(n-1)

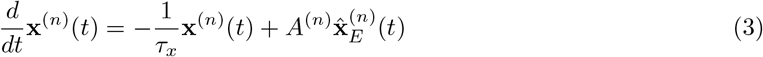

with *A*^(*n*)^ defined as an *M*-by-*M* matrix that implements a linear transformation of neural signals between nodes L(n-1) and L(n). The function of information transmission is attributed to E neurons (hence population readout in Eq. 3 is from E neurons), because long-range synapses between sensory brain areas are known to be excitatory ^27,20^. Specific signal transformations are aimed at creating layer-specific internal representations that are most useful for forming animal’s perception and for guiding behavior.

We next developed a biologically plausible spiking network for the generic processing layer L(n), *n* = 1, 2, …. Beyond L0, where efficient coding was performed by every single neuron, we now assume that the coding unit in L(n) is a population of recurrently connected 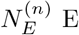 and 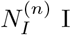 neurons, thus implementing a population code ^28,29,30^. Another crucial difference with L0 is that single neurons in recurrent layers receive converging inputs from several receptor types enabling the encoding of several variables simultaneously (i.e., mixed selectivity) ^31,32^, which has been suggested by empirical studies of neural responses in the cortex ^33,34,35^. Mixed selectivity of single neurons leads to simultaneous processing of multiple FF-driven features as well as to high-dimensional representations.

To accurately encode several stimulus features with a single population of neurons, we assumed that the loss function of each recurrent layers minimizes the encoding error across all *M* dimensions of the Targets 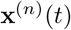 and their estimates 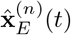 and 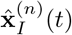. Same as in L0, the loss function of L(n) also includes the metabolic cost on spiking, with quadratic dependency on the spiking frequency. Because the encoding objective is now defined at the population level, the metabolic cost is considering the sum of neural activity across the population:

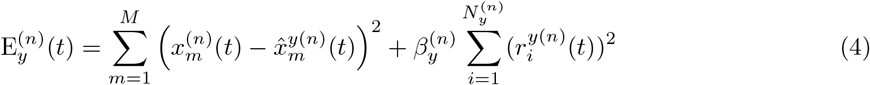

with constants 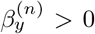 weighting the importance of the metabolic cost over the encoding error, and where 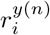 is a low-pass filtered spike train.

Assuming that a spike of a neuron occurs only if this decreases the instantaneous loss function of its population in Eq. 4 and adding noise for biological plausibility, we analytically derived a biologically plausible approximation of efficiency objectives in terms of a recurrently connected E-I spiking network with generalized LIF neurons (see Appendix A.4):

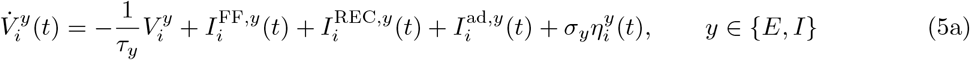

with fire-and-reset rule: if 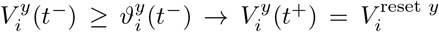. Note that we omitted the layer index (*n*) for intelligibility. Last terms on the right-hand side implement white noise in the membrane potential, where 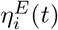 and 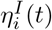 uncorrelated *i*.*i*.*d*. Gaussian variables with zero mean and unit variance. The dynamics of the membrane potential in Eq. 5a depends on spike-triggered adaptation currents and recurrent E-I synaptic currents that are detailed in the Appendix (A1.2), as well as on FF synaptic currents:

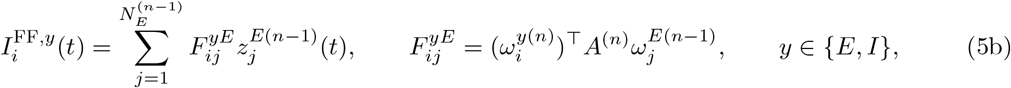

with 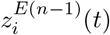 the synaptic (low-pass) filter of spikes of E neurons from layer L(n-1). The FF synapses in Eq. 5b depend on the similarity of the selectivity of the presynaptic neuron *j* and the postsynaptic neuron *i*, as well as on the transformation matrix *A*^(*n*)^. Matrix *A*^(*n*)^ implements a linear transformation of the population readout of E neurons from the previous layer L(n-1) (Eq. 3). The mechanistic effect of the transformation matrix *A*^(*n*)^ is therefore to shape the structure of the FF connectivity between processing layers L(n-1) and L(n).

### 2.2 Testing the efficient solution on a simplified model of sensory processing

To test our analytically derived model, we simulated a simplified model with 3 processing layers as on Fig. 1A, implementing neurons in layers 1 and 2 with the optimal solution in the Eq. 5a. Layer 1 received synaptic inputs from SA-LTMRs and RA-LTMRs in response to a naturalistic stimulus, modeled as Ornstein-Uhlenbeck process. We first assumed no transformation across layers (with *A*^(2)^ = **Id**) and identical parameters in L1 and L2, and we constrained parameters in L1 and L2 such that the simulated model minimized a weighted sum of the average encoding error and metabolic cost (see Appendix A.3). Optimal parameters of this simplified model were within biologically plausible ranges (SM Table S2) and were largely insensitive to the exact weighting of the error and the cost used to determine the optimum (Fig. S1-S3). This simple model reproduced several empirically measured parameter changes across layers of biological pathways, such as stronger I vs. E synapses ^36^, slightly faster membrane time constant in I compared to E neurons ^37,38^, synaptic time constants being faster than membrane time constants ^39^ and an increased ratio of E-to-I neurons in later ^40^ compared to earlier processing layers ^19^ (Fig. S2-3). This model had biologically plausible patterns of spiking activity (Fig. 1C) and it propagated fast and slow feature across processing layers with high accuracy (Fig. 1D, Fig. S4A). We measured the coefficients of determination for slow and fast feature of 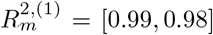 L1, and 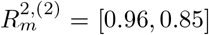 in L2. The mean firing rate was about 40 Hz in E and I neurons in L1 and E neurons in L2, and in was about 45 Hz in I neurons in L2 (Fig. S4B).

We tested the effect of removing the structure from the FF connectivity as prescribed by Eq. 5b, by randomly permuting the elements of FF connectivity matrices between L1 and L2 and distinguishing synapses to E (*F* ^*EE*^) and to I neurons (*F* ^*IE*^) in the postsynaptic layer. Random FF connectivity to E neurons caused a major increase in both the error and the cost, while random FF connectivity to I neurons increased the cost, but not the error (Fig. 1E). With random E-E connections, a random signal is transmitted from L1 to L2 that cannot be corrected by the structured signal received by I neurons, resulting in a strong increase in the encoding error. Meanwhile, the error increase with random FF E-I connectivity implies the transmission of a structured signal through FF E-E connections and a random signal through FF E-I connections. As I neurons are also driven by a structured signal through structured recurrent E-I connections in the local layer, random E-I FF connectivity is less disruptive to the representation in L2. However, random FF input to I neurons changes neural dynamics and increases the overall firing levels, thus causing a decrease in efficiency.

Next, we tested the benefit of across-type convergence of receptor neurons on efficient sensory processing in L1 and L2 by measuring the average encoding error and metabolic cost of networks with and without convergence. In networks without convergence, we set two unconnected E-I subnetworks with *N*_*E*_*/*2 E neurons per subnetwork, that only encoded a single FF variable each and otherwise had identical parameters as the network with convergence. We found that the networks without convergence had lower error but higher metabolic cost for both slow and fast features and in both recurrent layers (Fig. 2A). Across-type convergence of receptor neurons therefore benefits efficient transmission because it lowers the metabolic cost, with only a small increase in the encoding error. This benefit comes on top of that of within-type receptor convergence, which decreases the error on information transmission as reported above (Fig. 1B).

**Figure 2.**
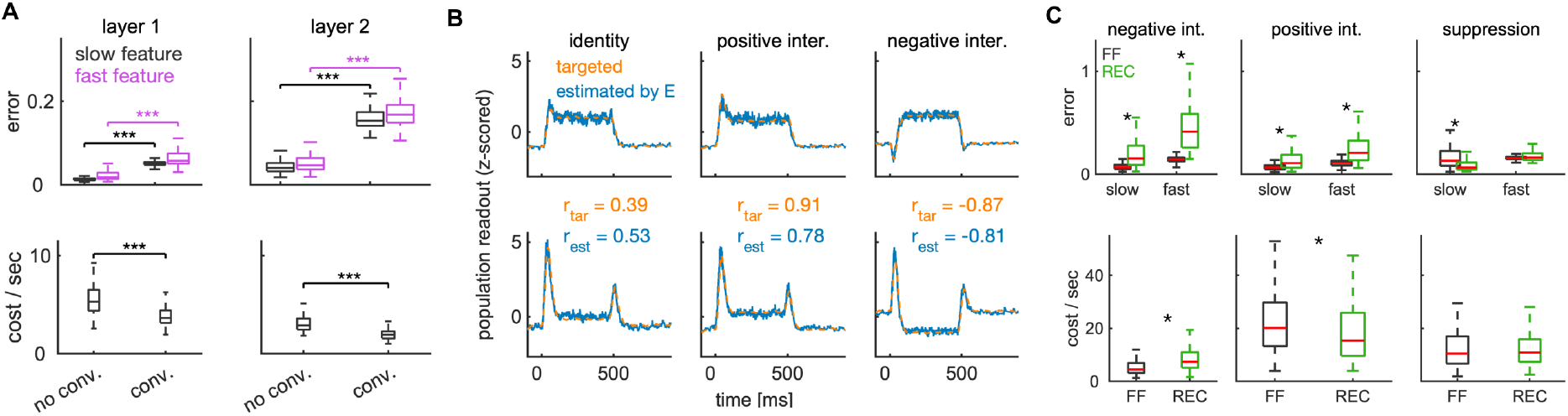
Efficient model has convergence of receptor neurons and can implement several types of feature mixing in the FF connectivity. **A** Error and cost per second for the efficient model (structured), and for models with randomly permuted E-to-E and E-to-I FF connectivity between L1-2. **B** Target and readout from E neurons in L2 of the slow (top) and fast (bottom) feature without transformation (left), positive interaction (middle) and negative interaction across features (right). We also report the Pearson’s correlation across features for the targets (in orange) and the estimates (in blue). **C** The error and the cost per second in models implementing a specific computation in the FF (black) and in the recurrent connectivity (green). Asterisk marks significance (*p <* 0.05, two-tailed t-test with Bonferroni correction). See SM Table S1–S2 for parameters.

Further, we tested the capability of the model to realize linear transformations on sensory features. We did so by modifying the transformation matrix *A*^(2)^ that determines the target signal in L2, which is mechanistically implemented by changing the structure of the FF connectivity between L1 and L2 (see Eq. 5b). For clarity of demonstration, we used a simpler stimulus, which was a step stimulus with duration of 500 milliseconds. As expected, in our model of receptor neurons, SA-LTMRs responded with increased firing during onset and sustained portions of the step and RA-LTMRs responded only at the onset and offset of the step (Fig. S4C). The model accurately realized a variety of signal transformations including positive and negative interaction of sensory features extracted by receptor neurons. We measured the coefficient of determination in L2 of *R*^2,(2)^ = [0.97, 0.95] for positive interaction, and *R*^2,(2)^ = [0.96, 0.94] for negative interaction, indicating excellent performance. A positive feature interaction makes interacting features more similar over time while negative interaction makes them more different, thus creating a temporal contrast between interacting features (Fig 2B). The interaction across features was implemented by setting the off-diagonal elements of the transformation matrix *A*^(2)^ to non-zero values. In particular, we manipulated off-diagonal elements that describe a symmetric interaction between the fast and slow feature, 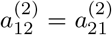. When these off-diagonal elements were set to a positive value (*a*_12_ = *a*_21_ = 0.6), the network realized a positive interaction between the slow and fast feature, making the readout of these two features more similar to each other (Fig. 2B, middle). We measured this effect by computing the Pearson’s correlation between the readout of these features. The correlation increased from 0.53 for the identity computation (*A*^(2)^ = **Id**) to 0.78 for the positive interaction. On the contrary, when the off-diagonal elements were set to a negative value (*a*_12_ = *a*_21_ = − 0.6), the network realized a negative interaction and a temporal contrast across features (Fig. 2B, right). Negative interactions led to anticorrelated population readouts, with Pearson’s correlation of −0.81. Similar results were obtained in I neurons (Fig. S4D).

Finally, we tested the capacity of the model to implement these population-wide computations though changes of the recurrent connectivity in L2 instead of the FF connectivity between L1 and L2. To implement the computations in L2, the FF connectivity between L1 and L2 was set to implement the identity computation and the recurrent E-E connectivity in L2 was modified to incorporate the matrix *A*^(2)^. This gave the recurrent synaptic weight between two E neurons in L2 according to the following rule: 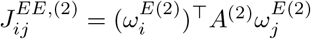. We found that negative interaction was more efficiently implemented in the FF connectivity, as it resulted in lower error in both features and in lower cost (Fig. 3A, left). Implementation of the positive interaction in FF compared to recurrent connectivity resulted in lower error but higher metabolic cost. Simpler computations that did not involve mixing across features were implemented with similar efficiency in FF and in recurrent connectivity. Suppression of the slow feature and simultaneous enhancement of the fast feature, achieved by manipulating the corresponding diagonal elements of the matrix *A*^(2)^, gave a slightly higher error on the slow feature when implemented in the FF connectivity, but the error on the fast feature and the metabolic cost were not significantly different. We thus conclude that linear and population-wide computations on sensory features in principle can be implemented by modulating the structure of recurrent instead of FF connectivity, however, the implementation of the negative interaction is more efficiently implemented in the FF connectivity.

**Figure 3.**
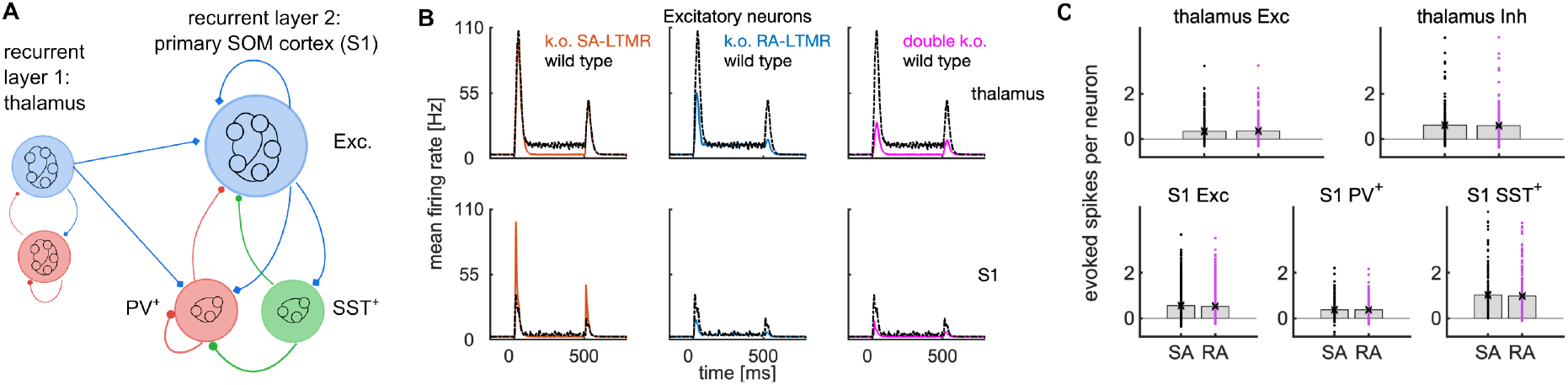
Model of information processing of the somatosensory system. **A** Schematic of L1 and L2 in a network modeling the somatosensory system. In L1 representing the thalamus, E neurons are unconnected. In L2 representing the primary somatosensory cortex, we have two types of I neurons. L0 is the same as on Fig. 1A. **B** Neuron and trial-averaged firing rate of E neurons in L1 (thalamus) and L2 (primary somatosensory cortex) in the receptor knock-out experiments. The knock-out is implemented as a complete removal (SA-LTMRs) and decrease of the readout of the activity of receptor neurons by the factor of 0.3 (RA-LTMRs), consistently with empirical study ^25^. **C** Trial-averaged number of spikes evoked in each single neuron in the thalamus (top) and in S1 (bottom) by 5 spikes of a single receptor neuron of type SA-LTMR or RA-LTMR. The distribution is across neurons and the bar marks the mean. In this figure, we use receptor neuron type-specific parameters in L0 and layer-specific parameters in L1-2 (see Tables S4–S5).

### 2.3 Nonlinear mixing of sensory features in mouse somatosensory cortex is explained by efficient multilayer networks with slow-fast temporal contrast encoding

We used our efficient multilayer network to capture information processing in a biological sensory pathway, the somatosensory system of the mouse. We used analytically derived efficient feedforward networks to capture receptor neurons as L0 and efficient recurrent E-I networks to capture the subcortical somatosensory system and the primary somatosensory cortex (S1) as L1 and L2, respectively. Analytically derived inhibitory neurons define like-to-like connectivity structure ^41,42^ and can be interpreted in a biological setting as parvalbumin-positive (PV) inhibitory neurons ^7^. In L2, we thus have E-PV, PV-PV and PV-E synapses ^43^ with like-to-like connectivity that implements lateral inhibition with a biologically plausible mechanism ^44^. Besides PV neurons, somatostatin-positive (SST) neurons are another type of I neuron that are important for signal processing in the cortex ^45^. We therefore added SST neurons to the S1 network and connected them with E and PV neurons using constraints reported in empirical studies of S1^38^. In particular, SST neurons received synaptic inputs from E neurons in the local layer, but they did not receive the FF synaptic inputs from the L1 as PV neurons did (Fig. 3A). Beyond these empirically observed constraints, we designed the E-SST and the SST-PV connectivity structure as like-to-like connectivity, while SST-E connectivity structure was designed as like-to-antilike, with synaptic weights negatively proportional to the similarity in encoding weights of the presynaptic SST and postsynaptic E neuron. Such like-to-antilike structure of SST-E connectivity implements winner-take-all inhibition ^46^ of E neurons.

Following our analytical results, the E-E synaptic connectivity has like-to-like structure, and its mean strength depends on the difference of time constants of the target and the population readouts (Appendix A.1.2). The empirical observation that subcortical somatosensory E neurons are largely unconnected was accounted for by assuming equal time constants between the target and the readout signals, because this set E-E connections in the subcortical system to 0. On the contrary, E neurons within S1 are known to be connected ^47,48,36^, and we captured this by setting the time constants of the target to be slower (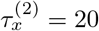 milliseconds (ms)) than the time constant of the readout (which is identical to the membrane time constant and was set to 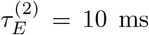). Sufficient strength of E-E synaptic connectivity in S1 was necessary to capture information processing of the mouse somatosensory system (Fig. S6). We stimulated our network with a step stimulus, for which the response profiles are known from empirical studies ^25,26^.

To capture the complexity of the mouse somatosensory system system, we allowed parameters of the model to differ between the two sensory neuron types and also between the L1 and L2 layers. We used biological parameters measured in the mammalian somatosensory system (see SM Table S5). In particular, we implemented a 2:1 ratio of the number of E to I neurons in L1^26^, while this ratio was 5:1 in L2^40^. The L2 network was much larger than the L1 network, with the total of 300 neurons in L1 and 1920 neurons in L2. Further, the mean synaptic connectivity PV-E (and E-PV) was stronger than mean SST-E (and E-SST) connectivity (see SM Table S6), which was important for capturing the information processing of the mouse somatosensory system (Fig. S6F).

Constrained with analytical results and biophysical parameters, our model reproduced several intriguing and so far unexplained observations in the mouse somatosensory system. We reproduced dynamical activity profiles observed in the thalamus and S1^25^, where average firing rates in the thalamus show significant sustained activity, while sustained activity is entirely suppressed in S1 (Fig. 3B, black).

Sustained activity is a dynamical feature of receptor neurons encoding the slow feature (Fig. S4C). The simplest hypothesis about why sustained activity is suppressed in S1 would be that the information about the slow feature propagates to the thalamus, but not to S1. To test this hypothesis, we performed perturbative experiments with weak signals, where we caused 5 spikes in a single receptor neuron and observed how such perturbation propagates across the 3-layer network. We found that such weak signals propagate well across the pathway, with similar effect of perturbation of SA-LTMRs and RA-LTMRs in both L1 and L2 (Fig. 3C). We found that the perturbation of a single SA-LTMR evoked on average 0.35 spikes in each E neuron in the L1 and 0.56 in L2, respectively, while single RA-LTMR evoked 0.36 spikes in E neurons in the L1 and 0.53 spikes in L2. Multiplying these numbers with the number of E neurons in each layer, we find that 5 spikes from a single SA-LTMR (RA-LTMR) evoked 70 (72) spikes in the L1 and 898 (848) in L2. This is about 14.2-fold amplification of the spiking signal from receptors to the L1, and 174.6-fold amplification between receptors and L2 (averaging across receptor types). These effects are in range of effects observed empirically in the mouse somatosensory system system ^25^. The perturbation of LTMRs also evoked a large number of spikes in I neurons in L2, with SST neurons showing stronger sensitivity than PV neurons (Fig. 3C, bottom middle and right, Fig. S5). Perturbation results clearly demonstrated the presence of the SA-LTMR signal in L2 and therefore did not support the hypothesis that a decrease of sustained activity between the L1 and L2 (Fig. 3B, black) occurs due to lack of propagation of SA-LTMR information to L2.

To get insight into processing of sensory features by the somatosensory system and further constrain our model, we next modeled the effect of knocking out by completely or partially disrupting ^24^ one or two types of receptor neurons. In real data, genetic knock-out of the RA-LTMR receptor neurons causes a decrease in firing rate response, while the knock-out of SA-LTMRs causes an increase at the onset and offset of the stimulus ^25^. We reasoned that such differential effects of the knock-out indicates a sublinear summation of the slow and fast feature in S1, and therefore a negative interaction between these features. We further hypothesized that suppression of sustained activity from thalamus and S1 might occur because of strong and precisely timed recurrent inhibition provided by PV and SST neurons on the activity of E neurons in S1. We tested these hypotheses by simulating the knock-out of receptor neurons. With an asymmetric negative interaction from slow to fast feature, our model reproduced activity profiles empirically observed in S1 with receptor knock-out (Fig. 3B, Fig. S5). We found a strong increase in sensitivity in L2 with k.o. of SA-LTMRs on both onset and offset responses, with ratio of peak firing rate in E neurons of 2.9 at the onset and of 3.0 at the offset. The same ratio with the k.o. of RA-LTMRs was 0.53 and 0.34 at the onset and offset, respectively. Both types of I neurons in S1 showed activity profiles as E neurons, with increased sensitivity with k.o. of SA-LTMRs and decreased sensitivity with k.o. of RA-LTMRs (Fig. S5).

In particular, with the removal of SST neurons, and thus the removal of winner-take-all type of inhibition, we still observed an overshoot of the firing rate with the k.o. of SA-LTMRs at the offset, but not at the onset. On the other hand, a strong decrease of connectivity with PV neurons abolished the overshoot at the onset and offset (Fig. S6).

An asymmetric negative interaction from the slow to the fast feature was necessary to recapitulate dynamical activity profiles in the somatosensory system, as no interaction, positive interaction, or symmetric negative interaction did not reproduce the differential effect on firing rates with the k.o. of SA-LTMRs and RA-LTMRs. Moreover, appropriate levels of recurrent inhibition by PV and SST neurons were both necessary to reproduce all empirically observed effects. We observed an overshoot of the firing rate with the k.o. of SA-LTMRs at the offset in some cases (no interaction, positive interaction and with disconnected SST neurons), however, a significant overshoot at the onset was not observed (see Fig. S6 for all). We therefore concluded that asymmetric negative interaction of fast and slow features is important for normal functioning of receptor neurons. Finally, sufficiently strong recurrent E-E synapses in L2 were also necessary to reproduce observed activity profiles.

## Discussion

We provided a theoretical framework for a mechanistic model of information encoding, transformation and transmission across a sensory pathway. At the entry level of sensory processing, our optimally efficient solution is given by feedforward networks of spiking neurons, modeling sensory receptors. Optimal transmission between receptors and the next processing layer is achieved with within-type and across-type convergence of sensory neurons on their postsynaptic targets in the next layer. In recurrent layers, optimally efficient solution is an E-I network of generalized integrate-and-fire neurons with mixed selectivity and structured recurrent connectivity. Optimally efficient transmission and signal transformation between two recurrently connected layers requires specific structure of the FF connectivity that at the same time enables efficient transmission and transformation of population-level signals. This specific structure depends on the type of computation performed by the network.

We found that within-type receptor convergence lowers the error of signal transmission, while across-type receptor convergence lowers the metabolic cost of encoding of multiple population-level signals by the same network in recurrently connected layers. Single receptor neurons only provided a noisy and imprecise estimate of the target, however, averaging the readout over many receptor neurons resulted in a very precise estimate of the target signal. We have shown that realizing computations by changes to the recurrent instead of FF connectivity is typically more efficient, in particular in case of a negative interaction across features. Our model demonstrated the relevance of highly time-sensitive and efficient neural code for neurobiology and suggested a computation of a time-dependent contrast (negative interaction) between slow and fast somatosensory features as computation performed in the somatosensory system. We also demonstrated the important role of strong recurrent inhibition with the temporal precision on the order of milliseconds, and suggested different computational roles for PV and SST types of inhibitory neurons for somatosensory processing.

While we here applied the model to processing of fine touch in the somatosensory system, our normative approach is general and can be easily adapted to study processing of other types of somatosensory stimuli, and potentially also to other sensory modalities.

### Limitations

Our model has several limitations. First, our model only describes an ascending sensory pathway and does not include feedback connectivity. While perception in awake animals is presumably mainly based on ascending processing ^49^, descending processing might be important in behavioral states such as drowsiness or sleep. External currents may be used to incorporate unstructured top-down feedback into efficient spiking networks ^44^. Including structured feedback, however, remains a challenge for future work. Second, we did not provide analytical solutions for SST neurons that provide winner-take-all inhibition in the last recurrent layer. Analytical derivation of such inhibition might require incorporation of computational objectives beyond encoding, transformation and transmission of sensory signals. Third, we only considered linear transformations at the network level. While the brain might also implement non-linear transformations, we showed that linear transformations reproduce effects in the somatosensory system of the mouse, which speaks to the relevance of linear transformations for neurobiology. Note that while signal transformations developed here are linear at the network level, they are highly non-linear at the single neuron level. This is demonstrated by the strongly non-linear effect of negative feature interaction on firing rates that are measured in single neurons.

## Acknowledgements

We acknowledge funding by NIH Brain Initiative (grants U19 NS107464, R01 NS109961, R01 NS108410 to SP), by Simons Foundation for Autism Research Initiative (SFARI; grant 982347 to SP) and a NIH Grant R00 NS119739 (to AJE).

## A Appendix / Supplementary material

### A.1 Analytical derivations of dynamical equations for efficient spiking models

#### A.1.1 Optimal encoding with a single sensory receptor neuron

In this section, we provide a detailed derivation of the spiking model in terms of a generalized leaky integrate-and-fire (gLIF) neuron with spike-triggered adaptation. Such model is derived from an assumption that the neural representation is encoded by a single neuron minimizing its encoding error and the metabolic cost on spiking. This model is relevant for modeling the activity of unconnected spiking neurons in the sensory periphery, such as mechanoreceptors in the somatosensory system.

We define a variable *f*_*i*_(*t*) as a series of uniform impulses (the spike train) emitted by a neuron *i*

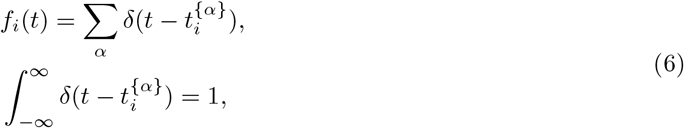

with 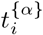 the spike time of the *α*−th spike of neuron *i*. Moreover, we define *z*_*i*_(*t*) and *r*_*i*_(*t*) as convolutions of the spike train *f*_*i*_(*t*) with an exponential low-pass filter,

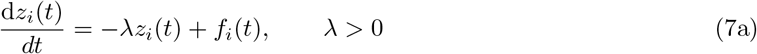

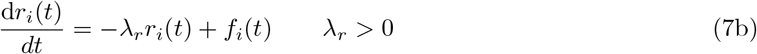

with inverse time constants *λ* and *λ*_*r*_, respectively.

##### Proposition 1.

*The instantaneous encoding error of a single spiking neuron minimizing a sum of quadratic encoding error and quadratic metabolic cost with every spike is bounded*.

*Given a continuous neural signal x*(*t*) ∈ ℝ ^[1*x*1]^ *and assuming the estimate of the signal by the neuron is* 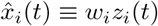, *with w*_*i*_ ∈ ℝ ^[1*x*1]^, *w*_*i*_ *>* 0 *the decoding weight of the neuron, we pose an objective function that minimizes the time-dependent, squared error between the signal x*(*t*) *and its estimate* 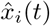, *and the metabolic cost incurred by neuron’s spiking activity:*

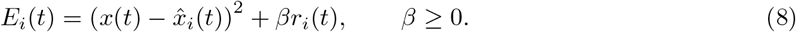

*Assuming that the spike is emitted if and only if it minimizes the objective function in the next time step, we get the following expression:*

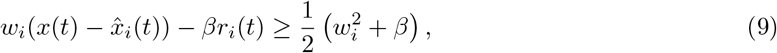

*and find that the encoding error is bounded as follows:*

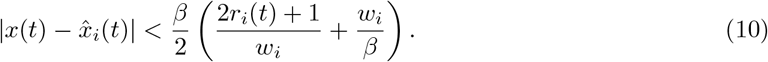

*With finite parameters β and w*_*i*_ *and a finite firing frequency of the neuron* 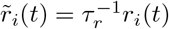, *the instantaneous encoding error by the neuron i*, 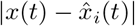 *is finite*.

*Proof*. Let *t*^−^ *< t* = *t*^spike^ *< t*^+^, with *t*^−^ the time just before a spike, *t*^spike^ the time in which a spike occurs, and *t*^+^ the time immediately after a spike. Following the definition of low-pass filtered spike train *r*_*i*_(*t*) (Eq. 7b) and of the estimate 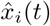, the effect of a spike at time *t* = *t*^spike^ is:

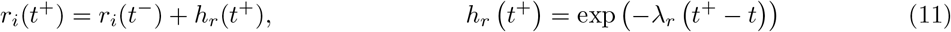

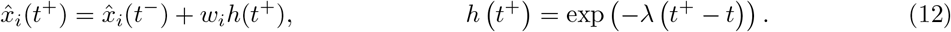

The assumption that a spike is fired at time *t*^+^ only if this decreases the error function is developed as follows:

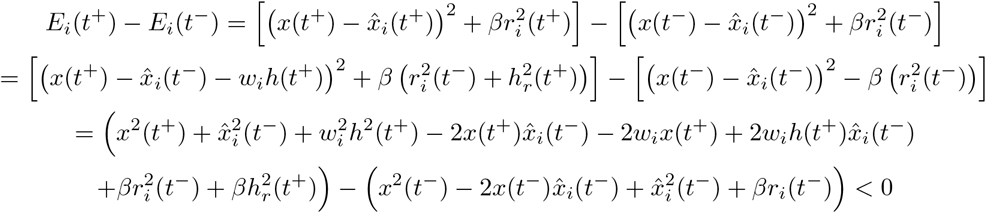

Taking one-sided limits from below, *t*^−^↑ *t*, and from above, *t*^+^ ↓ *t*, and since *x*(*t*) is a continuous function with *x*(*t*^−^) = *x*(*t*) = *x*(*t*^+^), we can simplify a number of terms, and get:

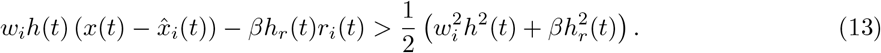

Using the continuity of the exponential function, we obtain

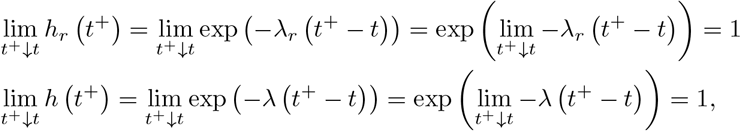

in the limit of *t*^+^ ↓ *t*. We can therefore simplify the Eq. 13 to get the Eq. 9.

From the Eq. 9 we interpret the left-hand side as the membrane potential of the neuron *i* and the right-hand side as the firing threshold. We therefore define the following variables:

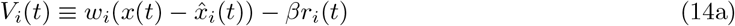

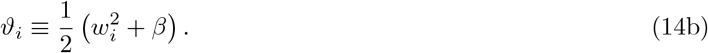

Moreover, we define the target signal *x*(*t*) through

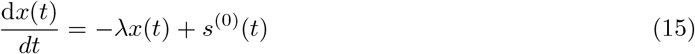

with *s*^(0)^(*t*) a continuous function. This leads to the following proposition.

##### Proposition 2.

*A spiking neuron that minimizes the sum of instantaneous quadratic encoding error and quadratic metabolic cost with every spike is a leaky integrate-and-fire neuron with spike-triggered adaptation*.

*Using the definition of the membrane potential, firing threshold and the target representation as in Eq. 14a-15 and defining the estimate of the target representation as* 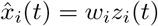 *with the convolution of the spike train z*_*i*_(*t*) *as in Eq. 7a, we find that the encoding neuron is an integrate-and-fire neuron with adaptation, obeying the following update equation:*

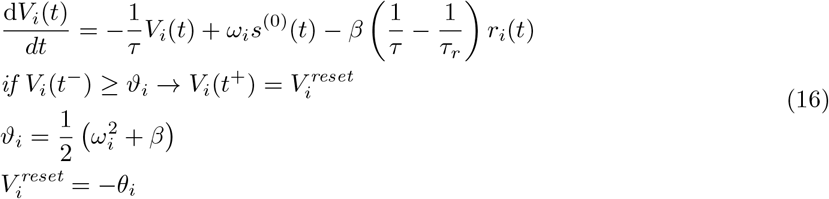

*with time constants τ* = *λ*^−1^ *and* 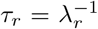 *and with constraints on selectivity weights: w*_*i*_ ≥ 0, *on the feedforward stimulus s*^(0)^(*t*) ≥ 0 ∀*t and on time constants: τ*_*r*_ ≥ *τ*.

*Proof*. We evaluate the time derivative of the membrane potential, by taking the time-derivative of time-dependent terms in Eq. 14a,

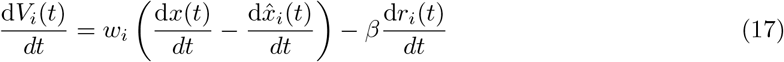

We use the definitions in Eq. 15 and Eq. 7b for the time-derivatives of the *x*(*t*) and *r*_*i*_(*t*), respectively, while the time-derivative of the estimate is

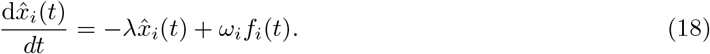

due to the expression for *z*_*i*_(*t*) in Eq. 7a and since we have that 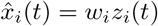. Using the time-derivatives of *x*(*t*), 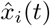 and *r*_*i*_(*t*), we get the following expression:

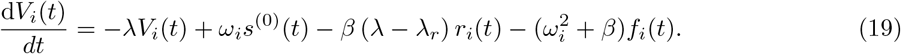

We now note that the last term on the right-hand side implements an instantaneous jump of the membrane potential upon the spike of the neuron. The instantaneous jump at the spike time is necessarily negative due to *β >* 0 (Eq. 8) and *r*_*i*_(*t*) *>* 0 ∀ *t* (Eq. 7b), which allows to interpret it as the reset current of an integrate-and-fire neuron. Applying the resetting of an integrate-and-fire neuron, we get the update equation for a spiking neuron in Eq. 16, and the reset current is given by:

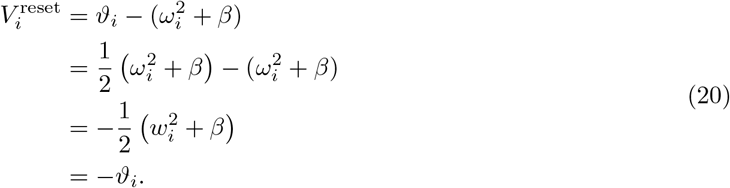

Following empirical evidence ^22,24^, receptor neurons always respond to the presence, and not to the absence of their preferred sensory feature, which constraints selectivity weights to positive values, *ω*_*i*_ *>* 0 ∀*i*. Stimulation of receptor neurons always causes a positive inward current ^25^, implying that the term *ω*_*i*_*s*^(0)^(*t*) is always positive, and leading to another constraint: *s*^(0)^(*t*) ≥ 0 ∀*t*.

Empirical studies of the somatosensory system further suggest the presence of adaptation in sensory mechanoreceptor neurons ^23^. Adaptation can be modeled as a hyperpolarizing current (negative with respect to the resting potential) in a LIF model, as its effect is to reduce the spiking frequency of the neuron during a short time after firing a spike ^50^. To interpret the third term on the right-hand side of Eq. 19 as an adaptation current, and since *β* ≥ 0 (see Eq. 8), we impose the following constraint on time constants *τ*_*r*_ ≥ *τ*.

#### A.1.2 Optimal encoding of high-dimensional neural representations with a recurrently connected E-I spiking network in L(n)

In this section, we provide detailed proofs of the recurrent spiking network in terms of a generalized LIF model neuron, assuming that a high-dimensional neural representation is encoded by a network of E and I neurons, and assuming the network is minimizing the encoding error and the metabolic cost. This model is relevant for modeling local microcircuits in subcortical and cortical sensory areas of mammals.

Assuming a population of *N*_*E*_ excitatory (E) and *N*_*I*_ inhibitory (I) neurons, we define convolutions of spike trains with an exponential filter,

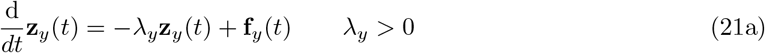

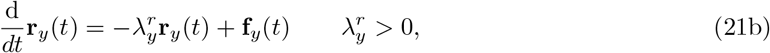

with **f**_*y*_(*t*), *y* ∈ *{E, I}* vectors of spike trains, 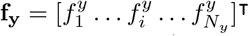. The spike train 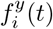 follows the definition in Eq. 6 and 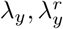 are inverse time constants, 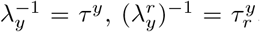. We further assume a target representation **x**(*t*) ∈ ℝ ^[*Mx*1]^, with *M* the maximal number of independent variables that can be encoded by the network, and define the estimate of the target representation as a linear sum of convolved spike trains across neurons,

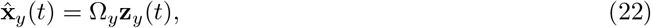

with 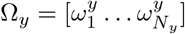 an [*M × N*_*y*_] matrix of decoding weights, and its columns 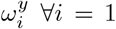, …, *N*_*y*_ are the decoding vectors associated to the neuron *i*. Using these definitions, we come to the following proposition.

##### Proposition 3.

*The instantaneous encoding error of a spiking network minimizing the sum of a quadratic encoding error and a quadratic metabolic cost with every spike is bounded*.

*We pose that the objective of the population y* ∈ *{E, I} is to minimize a quadratic encoding error between the target and the estimated signals* **x**(*t*) *and* 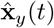 *and a quadratic metabolic cost on spiking activity that is proportional to the sum of the firing rates across neurons*,

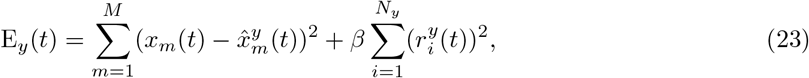

*with β >* 0 *and* 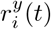 *the convolution of the spike train of the neuron i from Eq. 21b. Assuming that a spike of the neuron k at time t*^*spike*^ *is fired if and only if this decreases the objective function in the next time step, we find the following condition for threshold crossing of the neuron k:*

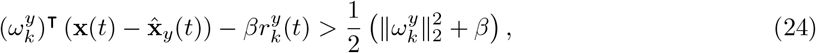

*with* 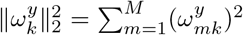 *the squared length of decoding vector of the spiking neuron. It follows that for any neuron i, the dot product of the encoding error and neuron’s decoding vector is bounded:*

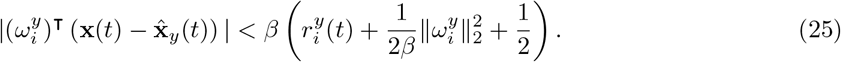

*With finite length of decoding vector* 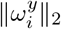, *finite metabolic parameter β and finite firing rate of the*

*neuron* 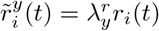, *the instantaneous encoding error of neuron* 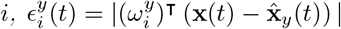, is *finite* ∀*i* = 1, …, *N*_*y*_.

*Proof*. We omit the population index *y* ∈ *{E, I}* for simpler notation, since the proof is the same for *y* = *E* and *y* = *I*. Let us have *t*^−^ *< t* = *t*^spike^ *< t*^+^, with *t*^−^ the time just before a spike, *t*^spike^ the time in which a spike occurs, and *t*^+^ the time immediately after a spike. The effect of the spike of the neuron *k* at time *t*^+^ on the convolved spike train *r*_*k*_(*t*) and on the estimate 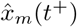 is, respectively,

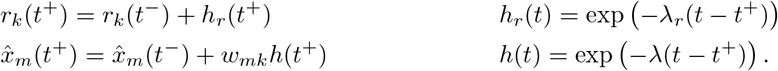

Using the assumption that a spike by the neuron *k* at time *t* = *t*^spike^ is fired only if it decreases the objective function,

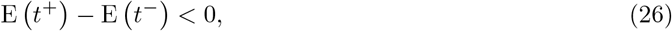

we evaluate the following:

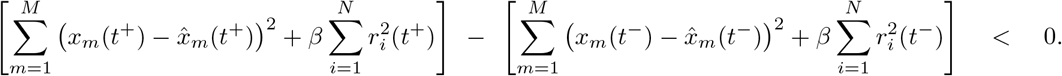

Inserting the effect of the spike on the convolved spike train *r*_*k*_(*t*) and on the estimate 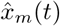, we get:

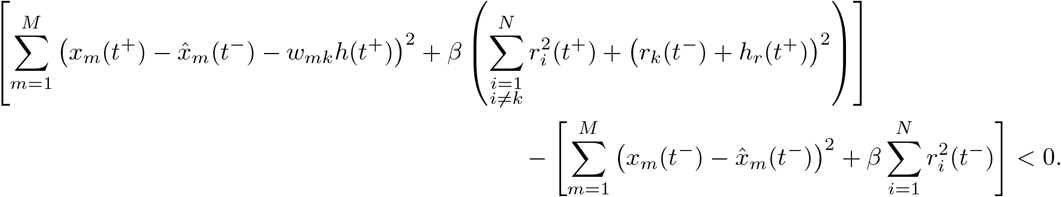

As we develop the squares, we get the following:

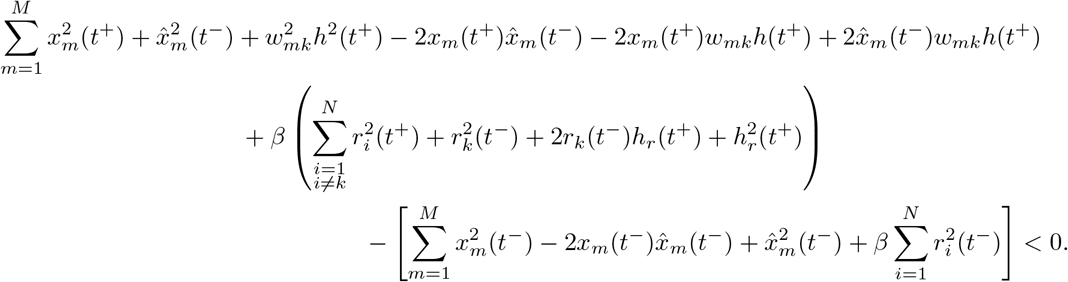

Taking the one-sided limit from below, *t*^−^ ↑ *t*, and from above, *t*^+^ ↓ *t*, we can simplify a number of terms and get the following exact solution:

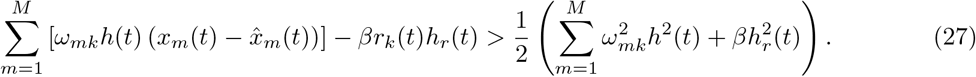

As we now evaluate the exact solution only for the time shortly after the spike, when *h*_*r*_(*t*) ≈ 1 and *h*(*t*) ≈ 1 (see section A.1.1), we get the following condition for spiking of the neuron *k*:

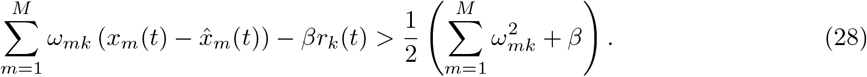

Rewriting the Eq. 28 in vector notation and adding the cell type index *y*, we get the Eq. 24 and the Eq. 25 follows immediately.

Using the condition for threshold crossing in Eq. 24, we define the membrane potential and the firing threshold of the neuron *i* of type *y* ∈ *{E, I}* as follows:

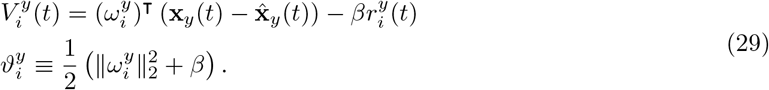

The target signal for the E and I population, meanwhile, is defined as

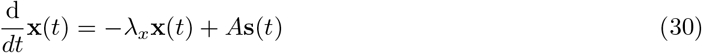

with **s**(*t*) a feedforward input to the network and *A* a linear transformation matrix. We naturally assume that the feedforward input to the network depends on the spiking activity of presynaptic neurons in an upstream (presynaptic) brain area. Following this assumption, we define **s**(*t*) as a weighted sum of low-pass filtered spike trains from a network of presynaptic neurons,

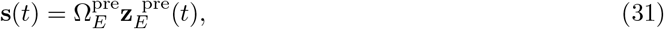

with 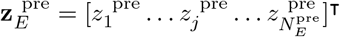 a vector of convolved spike trains from a network of 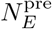 excitatory presynaptic neurons, and 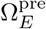 is the matrix of decoding weights of these neurons, with columns 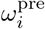 representing selectivity vectors of presynaptic neurons *i* = 1, …, 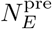. Moreover, we define a convolution of spikes with synaptic filters at E and I synapses *y* ∈ *{E, I}* as:

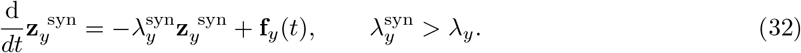

##### Proposition 4.

*A spiking network with excitatory and inhibitory neurons that minimizes a sum of instantaneous quadratic encoding error and quadratic metabolic cost with every spike is approximated as a network of generalized leaky integrate-and-fire neurons with structured connectivity*.

*Using the definitions of the membrane potential, the firing threshold and the target representation as in Eq. 29-31, the definition of estimates in Eq. 22 and the convolutions of spike trains in Eq. 21a and in Eq. 7b, a biologically plausible approximation of the optimal solution is an E-I network of generalized integrate-and-fire neurons with feedforward excitation, obeying the following update equations:*

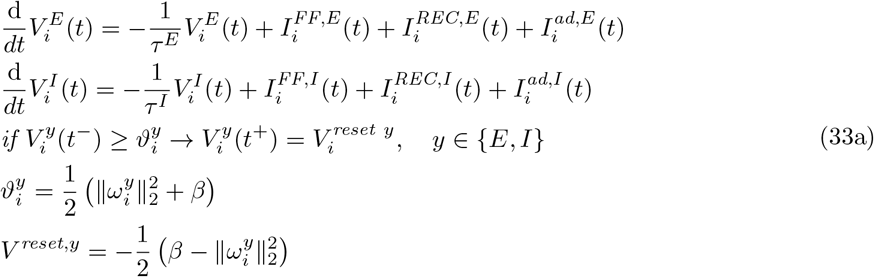

*with the following feedforward, recurrent and local currents:*

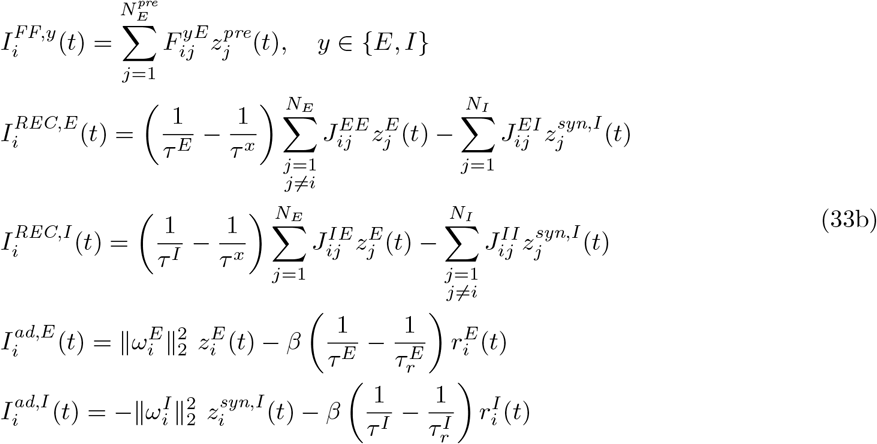

*with constraints on time constants: τ* ^*x*^ ≥ *τ* ^*y*^. *The synaptic weights between presynaptic neurons j and postsynaptic neurons i are as follows:*

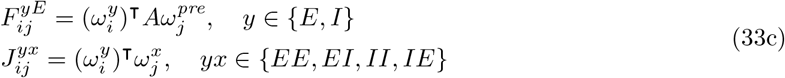

*with A positive semi-definite*.

*Proof*. We collect variables 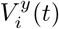 for *i* = 1, …, *N*_*y*_ across neurons of the same cell type into a vector, 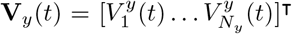 for *y* ∈ *{E, I}*. Similarly, we collect convolved spike trains across neurons and define a vector 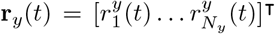. With these definitions, we can rewrite the membrane potential of *N*_*y*_ neurons for populations *y* = *E* and *y* = *I* in vector notation:

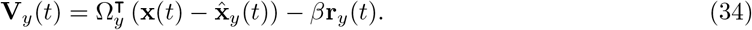

The time-derivative of **V**_*y*_(*t*) is then:

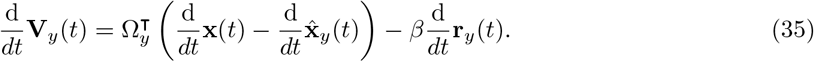

Using Eq. 30 and 31, we can write the time-derivative of the target signal:

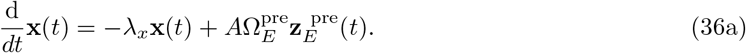

Moreover, from Eq. 22 and 32 we can write the time-derivative of the estimates of populations *y* ∈ *{E, I}*:

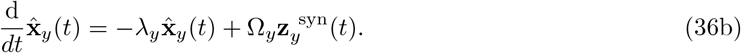

We first consider the case of E neurons. Inserting the time-derivatives from Eq. 36a, 36b and 21b into Eq. 35 for the case of E neurons (*y* = *E*), we get:

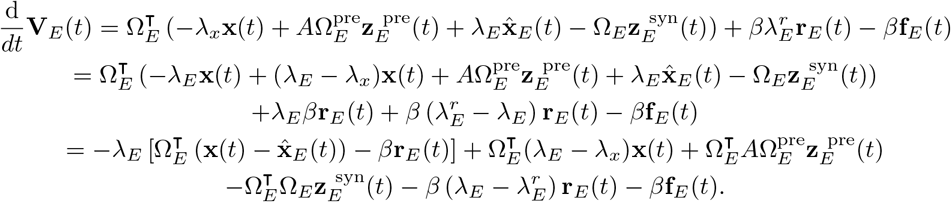

Noticing that the term in square parenthesis corresponds to the definition of the membrane potential of E neurons **V**_*E*_ in Eq. 34, we get the following exact expression in terms of a differential equation of the membrane potential of E neurons:

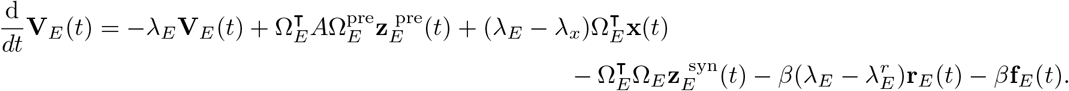

The third term on the right-hand side contains the target signal **x**(*t*), a quantity that is not directly computable by the network. Due to the minimization of the distance between the target signal **x**(*t*) and its estimate by E neurons 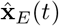 in the objective function in Eq. 23, we expect that the **x**(*t*) is well approximated by the 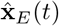, and make the following replacement: 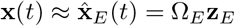.

Moreover, the fourth term on the right-hand side breaks Dale’s principle, since spikes of E neurons cause a negative current in postsynaptic neurons with similar selectivity. In this term and for pairs of E neurons *i* and *j* for which the following is true: 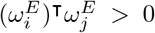, we have a negative synaptic current between E neurons. In biological networks, E neurons cannot generate inhibitory synaptic currents, and this term is therefore not biologically plausible. The term in question implements lateral inhibition between E neurons with similar selectivity ^10,9,4,11^, and it is natural to reason that in biological networks, inhibition is implemented by the activity of inhibitory neurons. To impose that the synaptic inhibition originates from the spiking activity of inhibitory neurons, we use the following approximation: 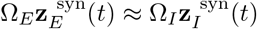. Applying these approximations, we obtain the following biologically plausible solution for E neurons:

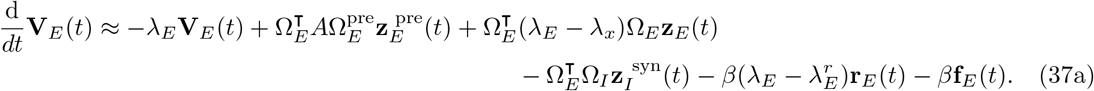

with constraints *λ*_*E*_ ≥ *λ*_*x*_, 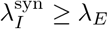, matrix *A* positive semi-definite, and non-negative elements of all decoding matrices, 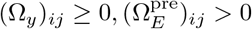.

We now consider the case of I neurons. We insert the time-derivatives from Eq. 36a, 36b and 21b into Eq. 35 for the case of I neurons (*y* = *I*), and get:

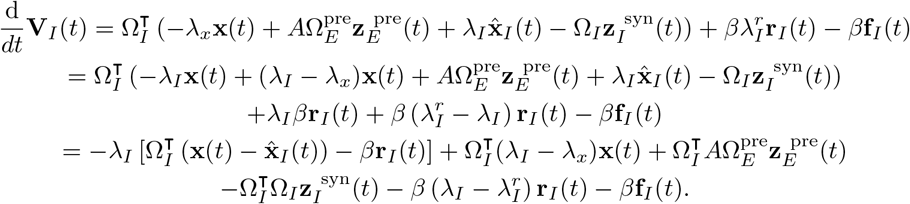

Similarly as in E neurons, the term in square parenthesis corresponds to the definition of the membrane potential of I neurons 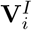 (Eq. 34 with *y* = *I*). Substituting **V**_*I*_ to the square parenthesis, we get the exact expression for the differential equation of the membrane potential of I neurons:

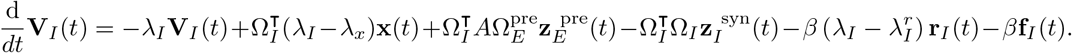

Similarly as in E neurons, the second term on the right-hand side contains **x**(*t*), a variable that we cannot safely assume to be directly computable through the network dynamics. We use the same approximation as in E neurons and substitute the target signal by its estimate, 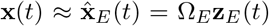. We obtain the following approximate expression for the membrane potential of I neurons:

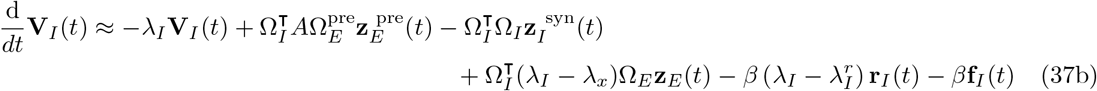

with constraints as in Eq. 37a, and, additionally, with constraint on inverse time constants *λ*_*I*_ *> λ*_*x*_, to ensure Dale’s principle in the fourth term on the right-hand side. Differential equations in Eq. 37a-37b are dynamical equations for the membrane potentials of E and I neurons, respectively.

Using time constants 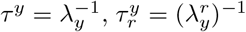 and 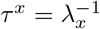, we now rewrite the Eq. 37a-37b for a single E and I neuron, respectively,

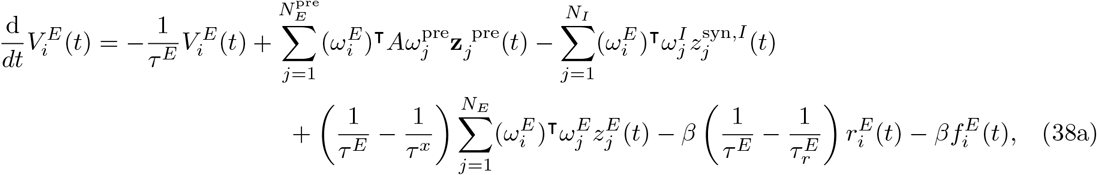

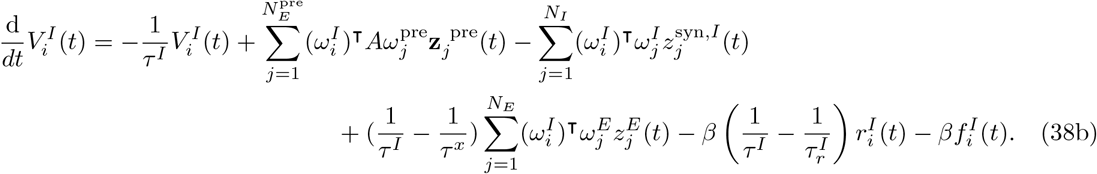

We note that last terms implement an instantaneous negative jump at the spike time of the neuron *i*. The instantaneous jump occurs as the neuron has reached the firing threshold and its effect is to reset the membrane potential immediately after a spike. This allows to omit the terms 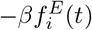 from Eq. 38a-38b and account for their their effect on the reset current *V* ^reset,*y*^,

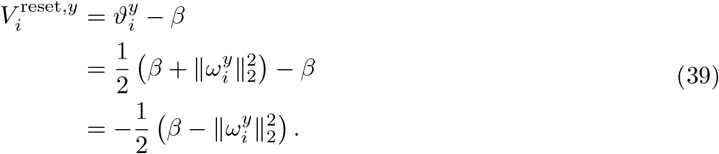

where we used the definition of the firing threshold 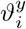 from Eq.29.

Moreover, to write the membrane equations in Eq. 38a-38b with a formulation that has a direct interpretation in neurobiology, we separated synaptic currents (between distinct neurons) from the self-generated currents (from the neuron to itself). In E neurons (Eq. 38a), a self-generated current that activates at the spike time is caused by the diagonal of the recurrent E-E connectivity,

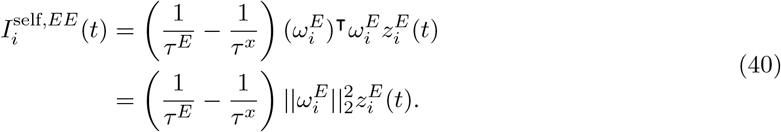

This term is always positive and has the dynamics of the low-pass filter 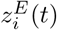. Similarly as in a related work ^7^, we interpret this self-excitatory current as a local depolarizing current. At a spike time of the E neuron *i*, the membrane potential 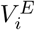 is reset (see Eq. 39), and the membrane potential is therefore strongly hyperpolarized. At the spike time activates also the local current 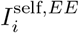, that rebounds the membrane potential of a hyperpolarized neuron back towards the firing threshold. This rebound current counteracts the hyperpolarization of the membrane potential after the reset.

Similarly, we have a self-generated current in I neurons, caused by the diagonal of the recurrent I-I connectivity in Eq. 38b,

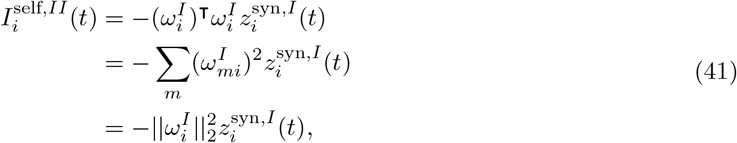

but in this case, the current is self-inhibitory. We interpret this current as a local hyperpolarizing current.

Taking into account the instantaneous resetting and the separation of self-generated and synaptic currents, we rewrite the membrane equation of a single E and I neuron as generalized LIF neurons:

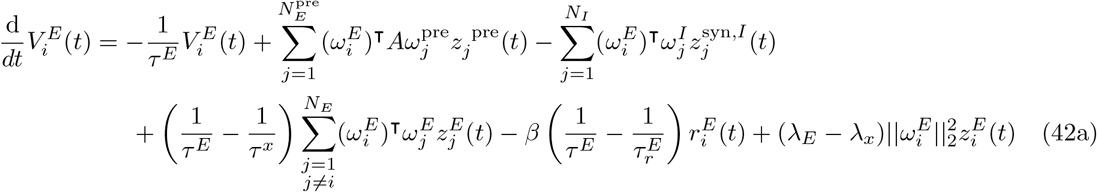

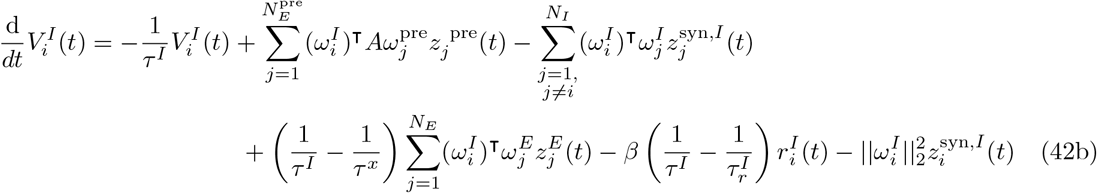

with fire-and-reset rule: if 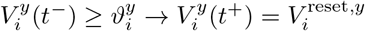, and the reset current 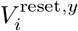 as in Eq. 39. The generalized LIF model has the following constraints: positive semi-definite transformation matrix *A, τ* ^*x*^ ≥ *τ* ^*y*^ and 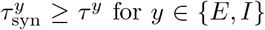 for *y* ∈ *{E, I}*. Constraints on time constants imply that the time constant of the target representation, *τ* ^*x*^, is equal or longer than the membrane time constant *τ* ^*y*^, and that the membrane time constant *τ* ^*y*^ is equal or longer than the synaptic time constant 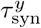.

By grouping currents in Eq. 42a-42b into feed-forward, recurrent and local currents, and defining the strength of feedforward and recurrent synapses as in Eq. 33c, we can rewrite the complete solution for an E-I spiking network as in Eq.33a, with currents as in Eq. 33b.

#### A.1.3 Optimal transmission between the receptor layer and the first recurrent layer

In the following, we develop the expression for the feedforward synaptic current from the receptor layer L0 to the first recurrently connected layer L1. We denote receptor neuron types with *m* = 1 …, *M* ^(0)^ and receptor neurons of the same type as 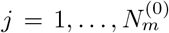, with (0) denoting the processing layer of receptor neurons. Moreover, we define the sensitivity of an E (I) neuron *i* in layer L1 to the activity of receptor neurons of type *m* as 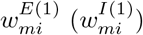. Following empirical observations in the somatosensory system, receptor neurons converge on neurons in the next processing layer ^22,23^. We distinguish across-type convergence, where receptors of several types have synapses with the same neurons in layer 1, and within-type convergence, where several receptor neurons of the same type have synapses on the same neurons in layer 1. Using the definition of sensitivity in layer L1 and taking into account within-type and across-type convergence of receptor neurons, we write the feedforward synaptic current to an E neuron *i* in L1 as follows:

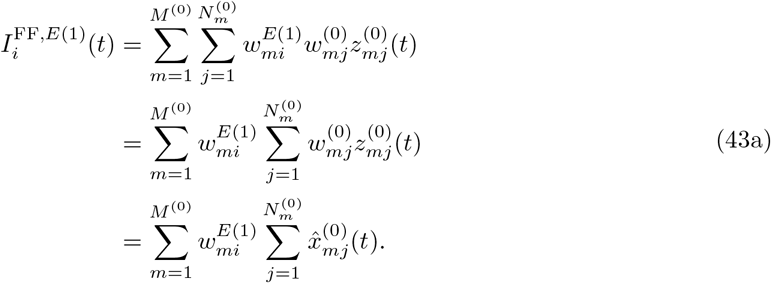

In the last line, we used the definition of the estimate 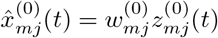. We get a similar expression for the feedforward current received by I neurons in layer L1:

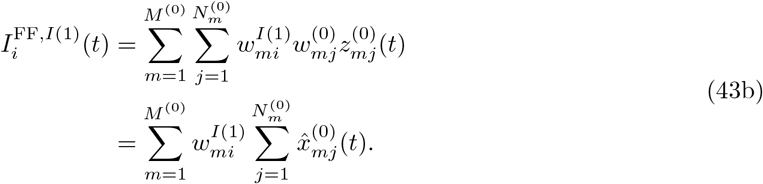

We now define the feedforward input relative to the stimulus feature *m*, 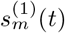, as the convergent readout from receptor neurons of type *m*,

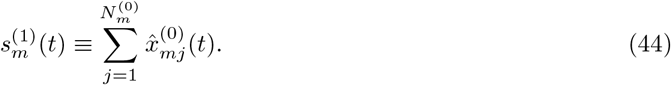

This allows to write the feedforward current to E and I neurons in L1 with a simpler expression:

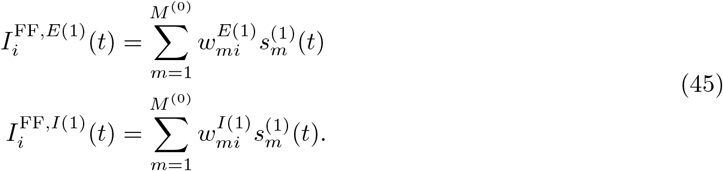

With Eq. 45, we recovered the expression for the feedforward current similar to the one used in the previous literature on efficient coding with spikes ^4,7^. We here demonstrated its biological plausibility by writing it as a sum of synaptic inputs from receptor neurons, which is a mechanistic model for the feedforward synaptic current in neurobiology.

We now rewrite the feedforward synaptic currents from L0 to L1 in Eq. 45 using synaptic connectivity matrices. From Eq. 43a-43b, we define synaptic connectivity matrices from receptor neurons of type *m* to E and I neurons in the L1:

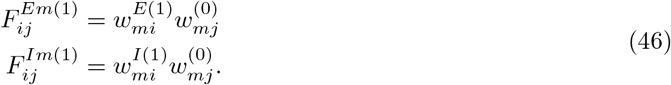

Following the empirical evidence in the somatosensory system of the mouse ^22,26^, we expect that the feedforward synapses from receptor neurons to E and I neurons in L1 are always excitatory. From Eq. 19, we have the constraint 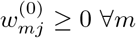, *j*. To guarantee excitatory effect of feedforward synapses between L0 and L1, we also have to constrain the selectivity weights in L1 to be non-negative, 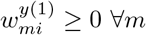, *i, y*. Using synaptic connectivity matrices, we can write the feedforward synaptic currents as follows:

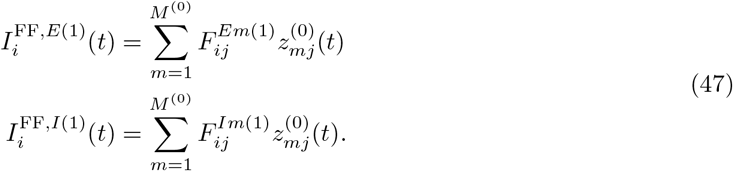

Expressions in Eq. 45 and Eq. 47 are equivalent.

### A.2 Computer simulations

All simulations were performed in Matlab (Mathworks), version 2022b and 2023b. To simulate the neural activity in receptor layer, we simulated differential equations for the membrane potentials 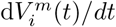 in Eq. 19 and low-pass filtered spike trains 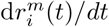 as in Eq. 7b. To simulate the neural activity in recurrent layers, we simulated differential equations for the membrane potentials d**V**_*y*_(*t*)*/dt, y* ∈ *{E, I}* using Eq. 37a and 37b. Moreover, we simulated low-pass filtered spike trains d**r**_*y*_(*t*)*/dt* using Eq. 21b and 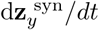 using Eq. 32, as well as estimates 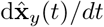 using Eq. 36b. In all layers, spike times were collected at the time of threshold crossing. All differential equations were simulated using the Euler method with time step of *dt* = 0.1 millisecond (ms). Initial conditions for the membrane potentials were drawn from a normal distribution, 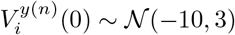, independently across neurons.

Previous work ^44^ derived a random Gaussian process (white noise) in the membrane potential of the spiking neuron from the assumption of a low-pass filtered noise in the condition for firing a spike. We added such noise term to our analytically derived equations for the membrane potential in all processing layers. Parameters of the Gaussian process are reported in Supplementary Tables (section A.7). The Gaussian process is independent across neurons.

We provided simulations of single trial activity to illustrate the examples of biologically plausible spiking dynamics. Furthermore, we iterated simulations across simulation trials whenever statistical measures and comparisons were performed. Iterating over simulation trials was necessary because or random initialization of the membrane potentials and because random variables that implement the noise in the membrane potential 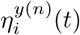 and selectivity weights 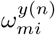 are particular instances of a random processes.

For results reported in section 2.3 (model of the somatosensory system), we only allowed the membrane potential noise 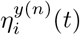 and the initial conditions to change across trials, while selectivity weights 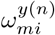 were drawn once at the beginning of the trial and were fixed across trials. This choice is justified by the fact that in a biological neural network, synaptic parameters do not change on short time scales. We did not notice a dependence of statistical results on a specific draw of selectivity weights in recurrent layers. However, in the model of the somatosensory system, some dependence was observed for parameters in the receptor layer where we used a very small number of receptor neurons to capture realistic neuron numbers stimulated by the step stimulus in empirical studies ^25^. These small neuron numbers implied very small sample sizes when drawing selectivity weights in L0.

### A.3 Performance measures

Measures of performance are based on the encoding error and the metabolic cost, since these quantities make part of the objective functions of the model (see Eq. 8 and 23). The encoding error is the mean squared error between the target signal *x*(*t*) and its estimate 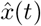, while the metabolic cost is proportional to the number of spikes fired by the network. In our model, using biologically relevant parameters and with neural firing rates in physiological ranges, the estimates 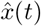 did not capture the mean and the variance of the target signal *x*(*t*), but they captured rather precisely the time-dependent fluctuations of the target signal, similarly as in a previous work on efficient coding ^12^. For this reason, we measured the encoding error on z-scored signals, with z-scoring operation defined as follows:

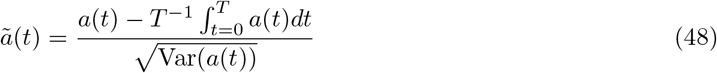

with *T* the length of the simulation trial and where the variance is computed over time.

To determine an optimal parameter across a range of tested parameters, we used a weighted sum of the encoding error and the metabolic cost. We also reported the performance of the network with coefficient of determination *R*^2^, a measure closely related to the mean squared error that has an intuitive interpretation in terms of units. In particular, the *R*^2^ = 1 indicates that the time-dependent fluctuations of the target signal are entirely predictable from the fluctuations of the estimate.

#### A.3.1 Performance measures in receptor layer (L0)

In receptor layer (L0), the time-dependent objective function is relative to the activity of single neurons (see Eq. 8), however, the readout of receptor neurons converges in L1 (see Eq. 43a-43b). Because of convergence, the effective estimate of the target signal is the sum across estimates of single neurons, 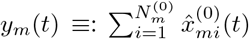, with *i* the neural index. We thus measure the accumulated representational error of L0 about the feature *m* as the average squared distance between the target signal 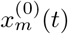 and the convergent readout *y*_*m*_(*t*):

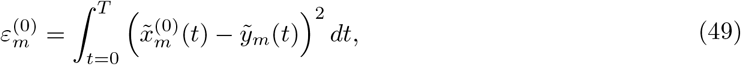

with 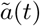 a z-scored signal as in Eq. 48. Note that all neurons of type *m* have the same target signal 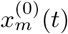.

The metabolic cost of receptor neurons of type *m* is:

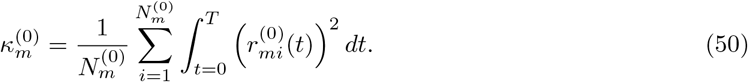

We used the encoding error of the convergent readout and the metabolic cost to define a heuristic measure of the average loss for the receptor type *m*:

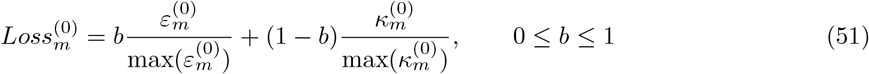

with max(*y*) taken across the range of values when exploring a specific parameter.

The coefficient of determination for the convergent readout of receptor neurons of type *m* is:

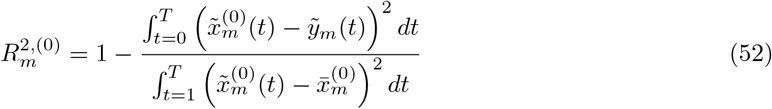

with 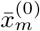 the average across time of 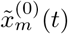. The coefficient of determination of 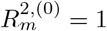 indicates that the fluctuations of the target signal 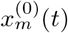 are entirely predictable from the fluctuations of the average convergent readout of receptor neurons of type *m, y*_*m*_(*t*).

#### A.3.2 Performance measures in recurrent layers

The encoding error in the processing layer L(n) about the feature *m* was measured as

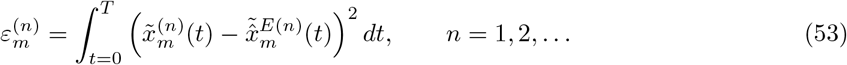

We denote with 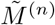 the number of features extracted from the external stimulus, and have 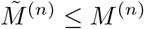. We averaged the encoding error across these feedforward-driven features,

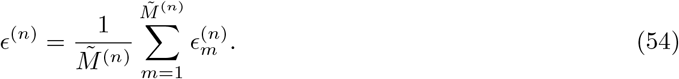

The metabolic cost in the processing layer L(n) was measured as:

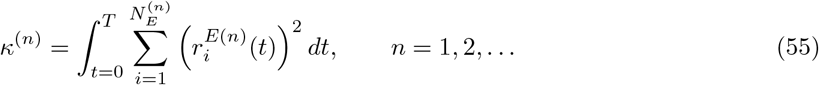

To determine an optimal parameter within a range of tested parameters, we used a heuristic measure for the average loss in layer L(n) defined as follows:

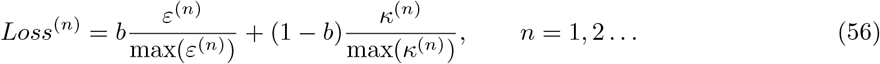

with 0 ≤ *b* ≤ 1 and with the max(*y*) operation taken over the range of results when exploring a specific parameter.

The coefficient of determination in the L(n) about the feature *m* is defined as:

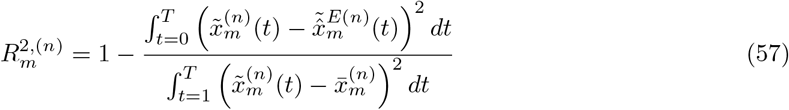

with 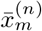 the average across time of 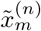. The coefficient of determination of 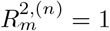 indicates that the fluctuations of 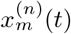 are entirely predictable from 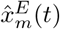.

### A.4 Parametrization of selectivity parameters

In the receptor layer, selectivity weights of 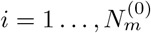 receptor neurons of type *m* were drawn from a normal distribution, 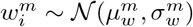, with 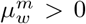. We used positive selectivity weights because mechanoreceptors were reported to only respond to the presence (and not the absence) of the features of tactile stimuli ^22^. In the recurrent layers, we draw selectivity of *i* = 1 …, *N*_*y*_ neurons to *m* = 1 …, *M* features from a normal distribution centered at zero, 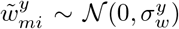. We rectified this distribution by setting all negative weights to 0, 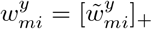 with [*x*]_+_ ≡ max(0, x) the rectified linear function. Parameters of distributions of selectivity weights are reported in Supplementary Tables (section A.7).

### A.5 External stimulus and stimulus features

We simulated the external stimulus *s*(*t*) as an Ornstein-Uhlenbeck process,

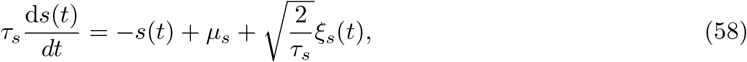

with parameters *τ*_*s*_ = 200 ms, *µ*_*s*_ = 50 *µN*, *σ*_*s*_ = 5 *µN* and with *ξ*_*s*_(*t*) a Gaussian noise with zero mean and covariance ⟨*s*(*t*_1_)*s*(*t*_2_) ⟩ = *δ*(*t*_1_ − *t*_2_). The Ornstein-Uhlenbeck process was chosen as an example of a biologically plausible stimulus, as we expect that a natural stimulus could be readily described by such a process.

Different types of receptor neurons *m* are sensitive to distinct features of the external stimulus. By studying the impulse response of slowly adapting and rapidly adapting low threshold mechanoreceptors (SA-LTMRs and RA-LTMRs, respectively), it has been shown that SA-LTMRs respond to the magnitude of the pressure applied to the skin, while RA-LTMRs respond to the time-difference in applied pressure, with an increase in the neural response to both positive and negative difference in the stimulus ^25^. Following these empirical measurements of the impulse response, we modeled the stimulus feature extracted by SA-LTMRs, 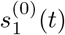, as proportional to the external stimulus, and the stimulus feature extracted by RA-LTMRs, 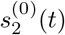, as proportional to the absolute value of the first temporal derivative of the stimulus:

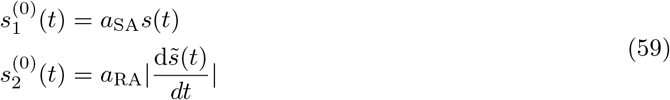

with 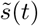 the convolution of *s*(*t*) with a normalized moving average (boxcar) filter of length 30 ms, to ensure the continuity of the signal. We set the constants *a*_SA_ and *a*_RA_ such that the firing rate averaged across time and across receptor neurons of the same type was approximately 25 Hz.

### A.6 Model parameters

#### A.6.1 Model parameters of the simplified model

To simulate the simplified model reported in Fig. 1-2, we used the set of parameters reported in Table S1, identical across receptor neuron types SA-LTMR and RA-LTMR. The neuronal activity in L1-2 was simulated with the set of parameters reported in Table S2, identical across L1 and L2. The biophysical parameters for L1 and L2 that are computed from model parameters on Table S2 are reported on Table S3. The adaptation strength in Table S3 is computed as: 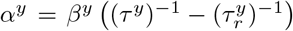, for *y* ∈ *{E, I}* (see Eq. 42a-42b).

#### A.6.2 Estimation of optimal parameters (simplified model)

Optimal parameters for the simplified model were estimated using the measure of average encoding error (Eq. 49 in L0 and Eq. 54 in L1-2), average metabolic cost (Eq. 50 in L0 and Eq. 55 in L1-2) and average loss (Eq. 51 in L0 and Eq. 56 in L1-2). The parameter search was iterated over simulation trials, because selectivity weights 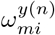 and the random variables that implement the noise in the membrane potential 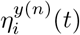 are particular instances of a random processes. We evaluated measures of the error and the cost in 150 simulation trials for each parameter, with one second of simulated time per trial. In every trial, we draw independently the selectivity weights, random variables that implement the membrane potential noise, and the initial conditions for the membrane potentials. Results were averaged across trials. Besides the parameter that is being tested, all other parameters were fixed as reported in Table S1 for the L0 and as in Table S2 for the L1-2. Optimal parameters are shown on Fig. S1 (L0) and Fig. S2-S3 (L1-2).

#### A.6.3 Model parameters that capture the somatosensory system

To capture the somatosensory system reported in Fig. 3, we used empirically observed parameters reported in the literature on neurobiology of the somatosensory system. We detailed a minimal set of model parameters necessary to reproduce our results in Table S4 (L0) and in Table S5 (L1 and L2). Table S6 reports biophysical parameters computed from model parameters in Table S5 and from model equations.

### A.7 Supplementary tables

**Table S1.** Table of model parameters in the L0 (simplified model).

| Symbol | Meaning | Value |
| --- | --- | --- |
| $M^{(0)}$ | number of receptor neuron types | 2 |
| $N_m^{(0)}$ | number of receptor neurons | 100 $\forall m$ |
| $\tau^m$ | membrane time constant | 14 ms $\forall m$ |
| $\tau_r^m$ | time constant of adaptation | 20 ms $\forall m$ |
| $\mu_w^m$ | mean of distribution of selectivity weights | 0.5 mV $\forall m$ |
| $\sigma_w^m$ | standard deviation of distribution of selectivity weights | 0.1 (mV)<sup>1/2</sup> $\forall m$ |
| $\sigma^m$ | noise intensity in the membrane potential | 0.7 (mV)<sup>1/2</sup> $\forall m$ |
| $\beta_m^{(0)}$ | metabolic constant | 12.5 mV $\forall m$ |

**Table S2.** Table of model parameters for generalized leaky IAF neurons in L1-2 (simplified model).

| Symbol | Meaning | Value |
| --- | --- | --- |
| $M$ | maximal number of represented variables | 5 |
| $N_E$ | number of E neurons | 400 |
| $N_E : N_I$ | ratio of E to I neurons | 2 |
| $\tau^x$ | time constant of the target signal $\mathbf{x}(t)$ | 20 ms |
| $\tau^E$ | membrane time constant E neurons | 10 ms |
| $\tau^I$ | membrane time constant I neurons | 8 ms |
| $\tau_r^E$ | time constant of adaptation E neurons | 10 ms |
| $\tau_r^I$ | time constant of adaptation I neurons | 10 ms |
| $\tau_{\text{syn}}^E$ | synaptic time constant of E synapses | 4 ms |
| $\tau_{\text{syn}}^I$ | synaptic time constant of I synapses | 4 ms |
| $\mu_w^y$ | mean of distribution of selectivity weights | 0 mV for $y \in \{E, I\}$ |
| $\sigma_w^E$ | standard deviation of distribution of selectivity weights in E neurons | 0.3 (mV)<sup>1/2</sup> |
| $\sigma_w^I : \sigma_w^E$ | ratio of standard deviation for selectivity weights I:E | 2.5:1 |
| $\sigma^E$ | noise intensity in the membrane potential of E neurons | 7.1 (mV)<sup>1/2</sup> |
| $\sigma^I$ | noise intensity in the membrane potential of I neurons | 8.0 (mV)<sup>1/2</sup> |
| $\beta^E$ | metabolic constant in E neurons | 36 mV |
| $\beta^I$ | metabolic constant in I neurons | 32 mV |

**Table S3.** Table of biophysical network properties of recurrent networks L1 and L2, calculated from model parameters in Table S2 (simplified model) and from model equations.

| Symbol | Meaning | Value |
| --- | --- | --- |
| $\langle F_{ij}^{EE} \rangle$ | mean synaptic strength feedforward E-to-E | 0.1 mV |
| $\langle F_{ij}^{IE} \rangle$ | mean synaptic strength feedforward E-to-I | 0.21 mV |
| $\langle J_{ij}^{IE} \rangle$ | mean synaptic strength recurrent E-to-I | 0.23 mV |
| $\langle J_{ij}^{II} \rangle$ | mean synaptic strength recurrent I-to-I | 0.6 mV |
| $\langle J_{ij}^{EI} \rangle$ | mean synaptic strength recurrent E-to-I | 0.23 mV |
| $\langle J_{ij}^{EE} \rangle$ | mean synaptic strength recurrent E-to-E | 0.1 mV |
| $p^{xy}$ | connection probability (all connectivity matrices) | $\sim 0.75$ |
| $\langle \vartheta_i^E \rangle_i$ | mean distance between threshold and resting potential in E | 18.1 mV |
| $\langle \vartheta_i^I \rangle_i$ | mean distance between threshold and resting potential in I | 16.5 mV |
| $\alpha^E$ | adaptation strength in E neurons | 0 mV |
| $\alpha^I$ | adaptation strength in I neurons | 0.9 mV |

**Table S4.** Table of model parameters for SA-LTMRs and RA-LTMRs (model of the somatosensory system).

| Meaning | Value SA-LTMR | Value RA-LTMR |
| --- | --- | --- |
| number of receptor neurons | 5 | 6 |
| membrane time constant | 10 ms | 15 ms |
| time constant of adaptation | 30 ms | 20 ms |
| mean of selectivity weights | 5 mV | 5 mV |
| standard deviation of selectivity weights | $3 \text{ (mV)}^{1/2}$ | $3 \text{ (mV)}^{1/2}$ |
| noise intensity in the membrane potential | $1 \text{ (mV)}^{1/2}$ | $1 \text{ (mV)}^{1/2}$ |
| metabolic constant | 120 mV | 120 mV |

**Table S5.** Table of model parameters for generalized leaky IAF neurons in L1 and L2 (model of the somatosensory system). Values for the SST neurons in L2 are equivalent to values for PV neurons, unless otherwise specified.

| Symbol (units) | Value in L1 | Value in L2 |
| --- | --- | --- |
| maximal number of represented variables | 5 | 5 |
| number of E neurons | 200 | 1600 |
| ratio of E to I neuron numbers | 2:1 | 5:1 |
| time constant of the target signal $\mathbf{x}(t)$ (ms) | 10 | 20 |
| membrane time constant of E neurons (ms) | 10 | 10 |
| membrane time constant of I neurons (ms) | 8 | 6 |
| time constant of adaptation in E neurons (ms) | 12 | 13 |
| time constant of adaptation in I neurons (ms) | 10 | 10 |
| synaptic time constant of E synapses (ms) | 4 | 4 |
| synaptic time constant of I synapses (ms) | 4 | 4 |
| shift of the mean of selectivity weights in dimensions [1,2] in E (mV) | [0,0] | [0,0] |
| shift of the mean of selectivity weights in dimensions [1,2] in I (mV) | [0.35,0] | [0,0] |
| standard deviation selectivity weights in E ($\text{(mV)}^{1/2}$) | 0.3 | 0.42 |
| ratio of standard deviations for selectivity weights I:E | 1.5:1 | [2.2:1,0.2:1] for PV, SST |
| noise intensity in the membrane potential of E neurons ($\text{(mV)}^{1/2}$) | 8.0 | 7.7 |
| noise intensity in the membrane potential of I neurons ($\text{(mV)}^{1/2}$) | 9.2 | 11.1 |
| metabolic constant in E neurons (mV) | 47.7 | 36.9 |
| metabolic constant in I neurons (mV) | 41.4 | 25.4 |
| synaptic delay in recurrent E-E synapses (ms) | 0 | 0.7 |
| synaptic delay in recurrent E-I and I-E synapses (ms) | 0 | 1 (E-PV), 0.5 (E-SST) |

**Table S6.** Table of biophysical network properties for the model of the somatosensory system, calculated from parameters in Table S5 and from model equations.

| Symbol (units) | Value in L1 | Value in L2 |
| --- | --- | --- |
| mean synaptic strength FF E-E (mV) | 0.1 | 0.12 |
| mean synaptic strength FF E-PV (mV) | 0.21 | 0.27 |
| mean synaptic strength REC E-PV (mV) | 0.16 | 0.40 |
| mean synaptic strength REC PV-PV (mV) | 0.35 | 0.94 |
| mean synaptic strength REC PV-E (mV) | 0.16 | 0.40 |
| mean synaptic strength REC E-E (mV) | 0.0 | 0.18 |
| mean synaptic strength REC E-SST (mV) | - | 0.04 |
| mean synaptic strength REC SST-PV (mV) | - | 0.09 |
| mean synaptic strength REC SST-E (mV) | - | 0.04 |
| connection probability FF to [E,I] | $\sim [0.75, 0.75]$ | [0.3,0.3] |
| mean distance between threshold and resting potential in E (mV) | 23.1 | 18.7 |
| mean distance between threshold to resting potential in I (mV) | 21.1 | 13.8 |
| adaptation strength in E (mV) | 0.80 | 0.85 |
| adaptation strength in I (mV) | 1.03 | 1.70 |

### A.8 Supplementary figures

**Figure S1.**
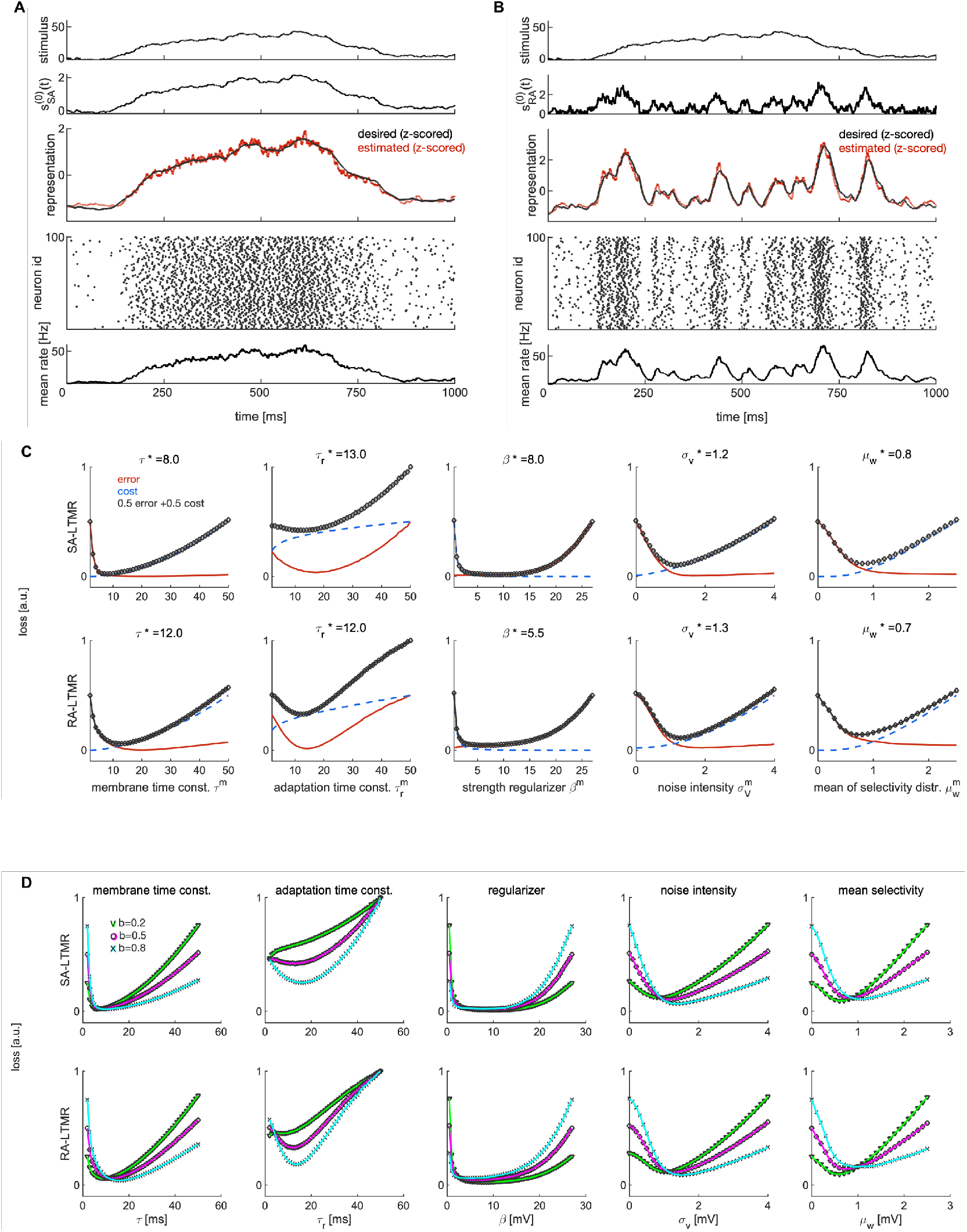
Activity and optimal model parameters in L0. **A** Activity of 100 receptor neurons of type SA-LTMR in a single simulation trial. Top: External stimulus is an Ornstein-Uhlenbeck process with the time constant of 200 ms. Second from top: Feature of the external stimulus extracted by SA-LTMR receptor neurons. Third from top: Target representation (black) and the population readout (red) of the spiking activity. Signals are z-scored since the model does not capture the absolute value, but time-dependent fluctuations of the target signal. Fourth from top: Spiking activity. Bottom: Firing rate, averaged across neurons. **B** As in A, but for RA-LTMRs. For the simulation in A and B, we used the same set of parameters, reported in Table S1. Accurate encoding of fast and slow stimulus features is therefore achieved for the same set of parameters. **C** The error (red), the cost (blue) and the average loss (black) as a function of model parameters in receptor neurons SA-LTMR (top) and RA-LTMR (bottom). The average loss is a weighted sum of the error and the cost, here reported for the equal weighting (b=0.5) For each plot, we vary a single parameter, while other parameters are as in Table S1. Above each plot, we mark with asterisk the parameter value that minimizes the average loss. **D** Same as in C, but showing the average loss for three different weightings of the error and the cost b.

**Figure S2.**
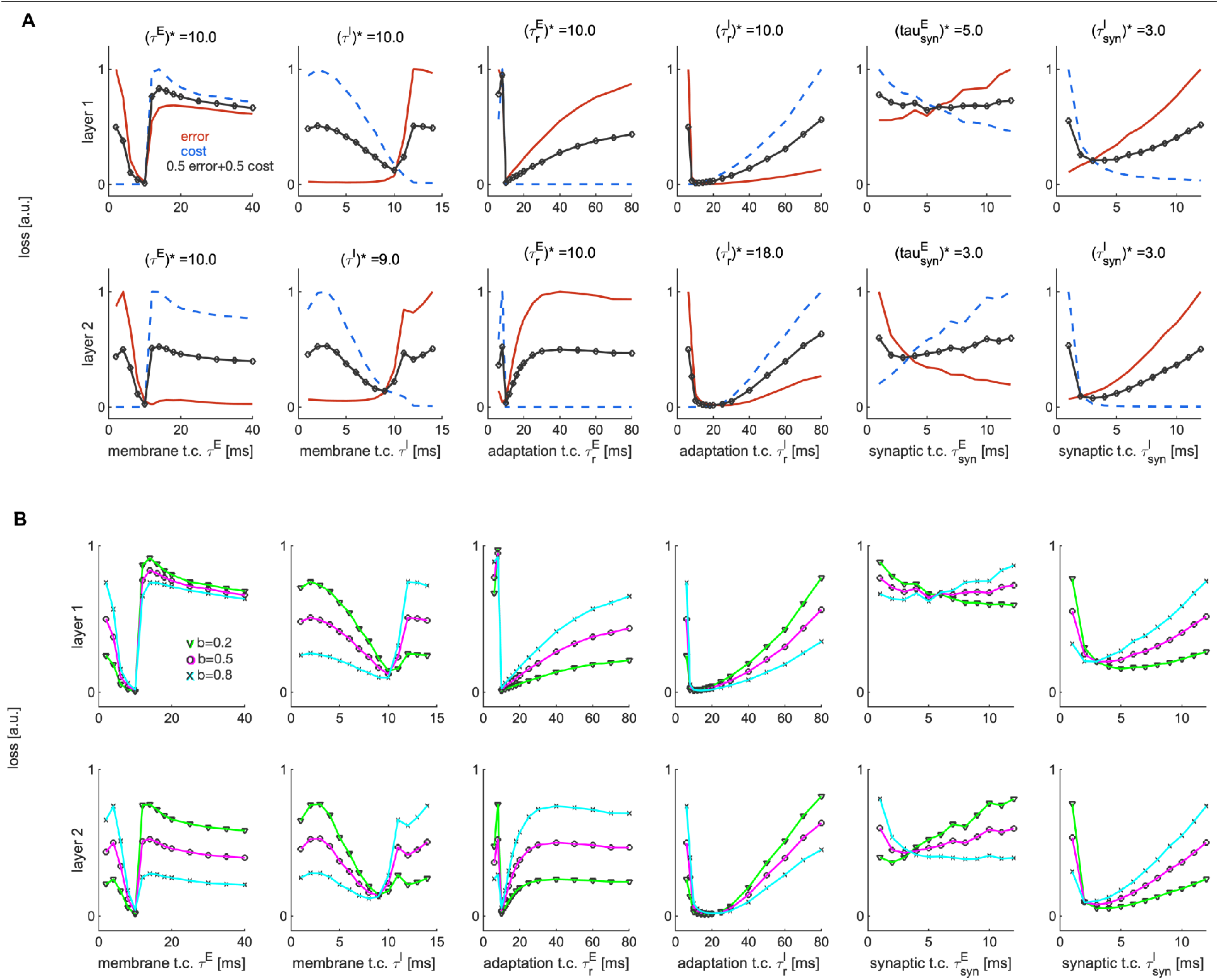
Optimal time constants in L1-2. **A** Same as in Figure S1C, but as a function of time constants in L1 and 2. The performance measures are measured separately for the L1 (top) and the L2 (bottom). L1-2 are simulated with the same set of parameters reported in Table S2. **B** Same as in A, showing the average loss for three different weightings of the error and the cost.

**Figure S3.**
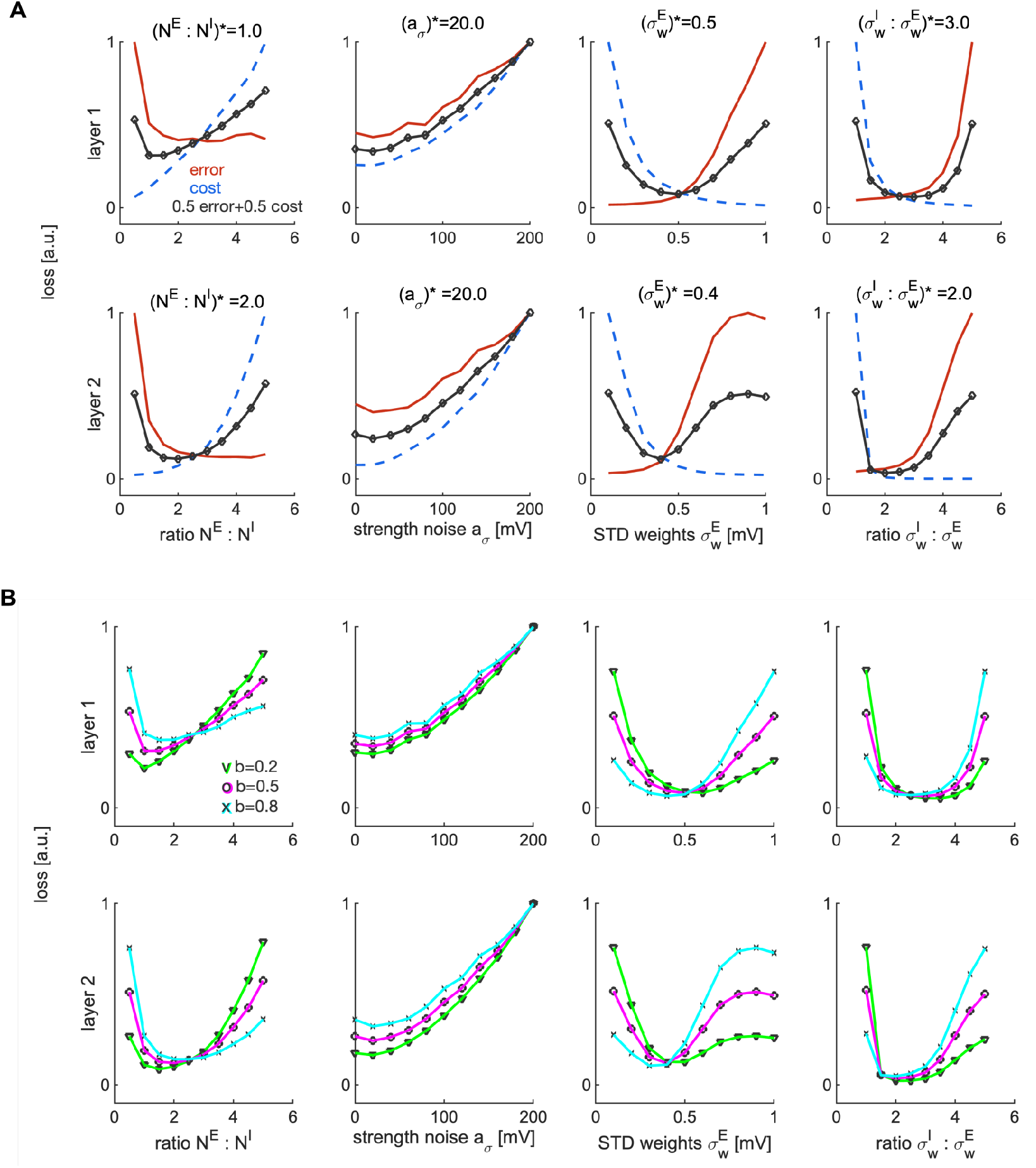
Optimal model parameters in L1-2. **A** Same as in Figure S1A, as a function of model parameters in L1 and 2. For the strength of the noise, we use a heuristic model with the noise intensity inversely proportional to the logarithm of the number of neurons, *σ*^*y*^ = *a*_*σ*_(log(*N* ^*y*^))^−1^. Since there are more E than I neurons, this gives stronger noise intensity and higher baseline firing rate in I compared to E neurons, as suggested by measurements of baseline firing rate in the primary somatosensory cortex ^25^ and in the visual cortex ^51^ of the mouse. **B** Same as in A, showing the average loss for three different weightings of the error and the cost.

**Figure S4.**
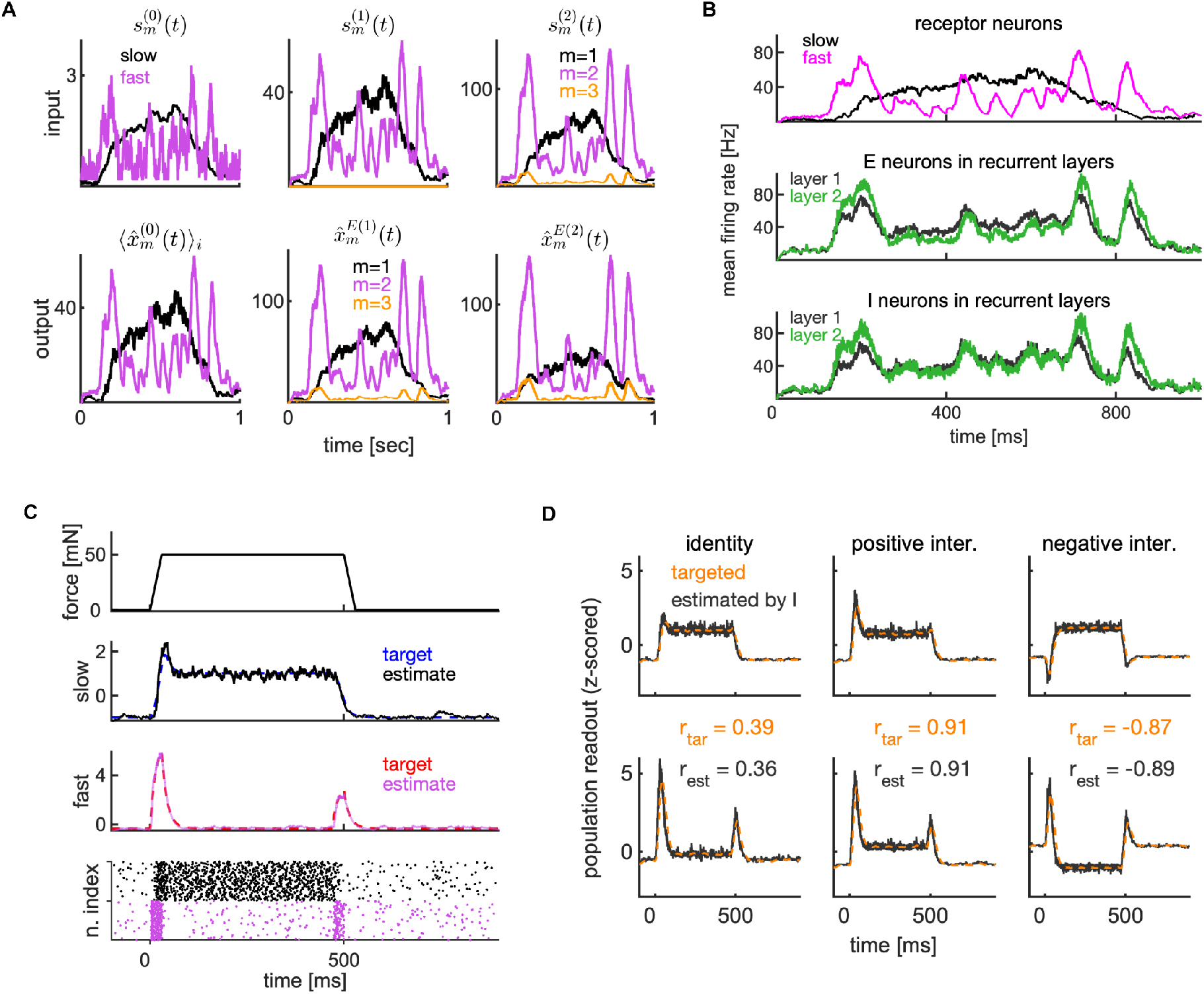
Efficient signal transmission across processing layers. **A** The FF input *s*^(*n*)^(*t*) (top) and the population readout of E neurons 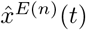 (bottom) across processing layers 0, 1 and 2 in a single simulation trial. By construction, the population readout of the previous layer is the FF input to the next layer, hence, the top middle plot is identical to bottom left, and the top right plot is identical to the bottom middle plot. **B** Firing rate averaged across neurons. **C** Top: External stimulus is a step stimulus with 30 ms ramp at the onset and the offset. 2nd and 3rd from top: Target and estimated representation of the slow and fast stimulus feature by SA-LTMRs and RA-LTMRs. Signals are z-scored. Bottom: Spiking activity of SA-LTMRs (black) and RA-LTMRs (magenta). **D** Target and estimate by I neurons in L2 of the slow (top) and fast (bottom) feature. We show the computation of identity function (left), positive interaction (middle) and negative interaction across features (right). We report the Pearson’s correlation across fast and slow feature for target signals (in orange) and for the estimates (in black). See Tables S1–S2 for parameters.

**Figure S5.**
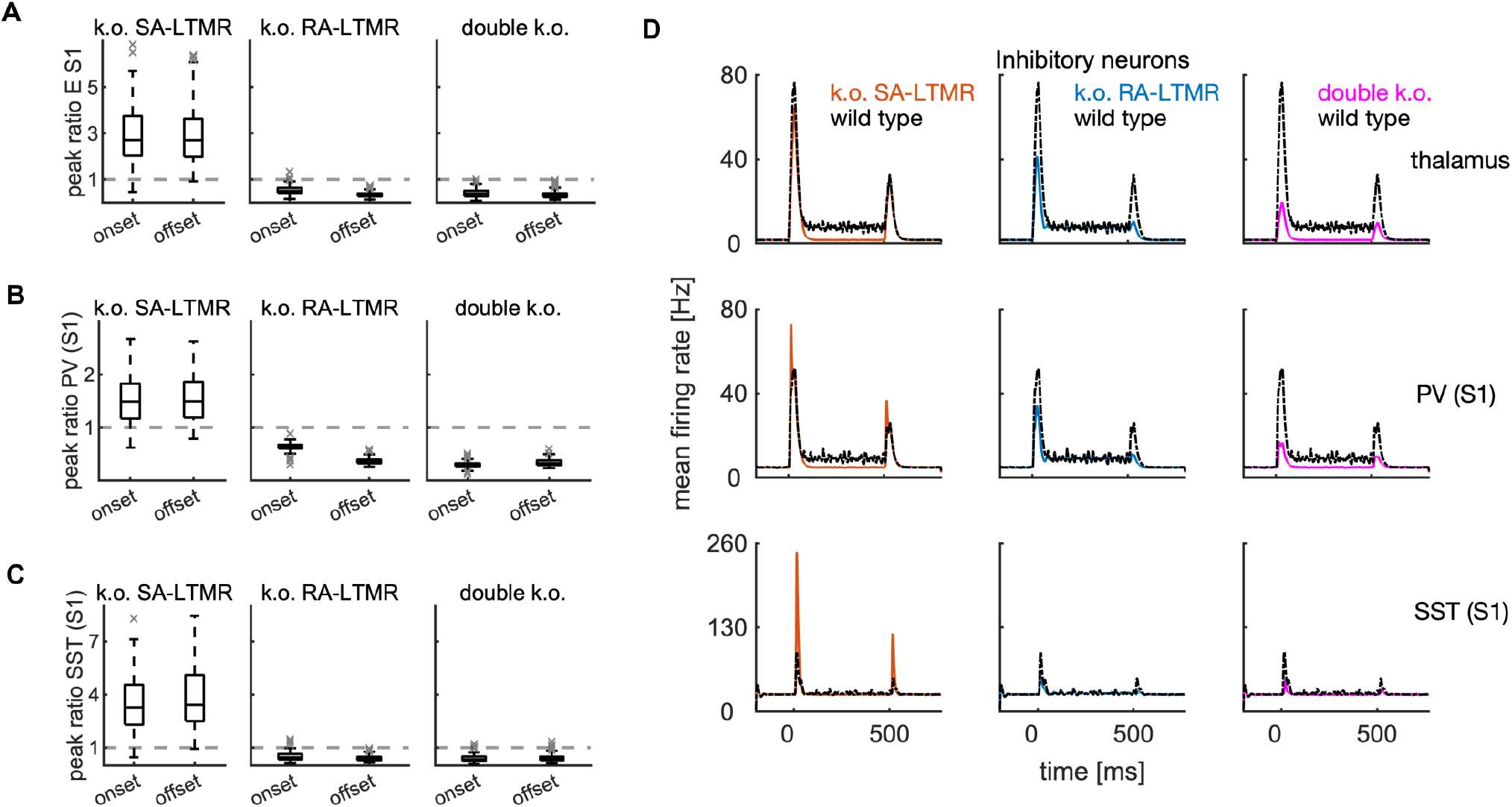
Effect of receptor knock-out on firing rates in the thalamus and S1. **A** Ratio of maximal firing rate with and without receptor knock-out in E neurons in S1. We show results for the knock-out (k.o.) of SA-LTMRs, RA-LTMRs and for the double knock-out of both receptor types. Maximal firing rates are measured in the time window of [0,50] ms with respect to the onset and offset of the step stimulus and are trial-and neuron-averaged. **B** Same as in A, showing the ratio of maximal firing rates in PV neurons. **C** Same as in A, showing the ratio of maximal firing rates in SST neurons. **D** Neuron-averaged firing rate in the thalamus and in PV and SST neurons in S1 in the knock-out conditions and in the wild type.

**Figure S6.**
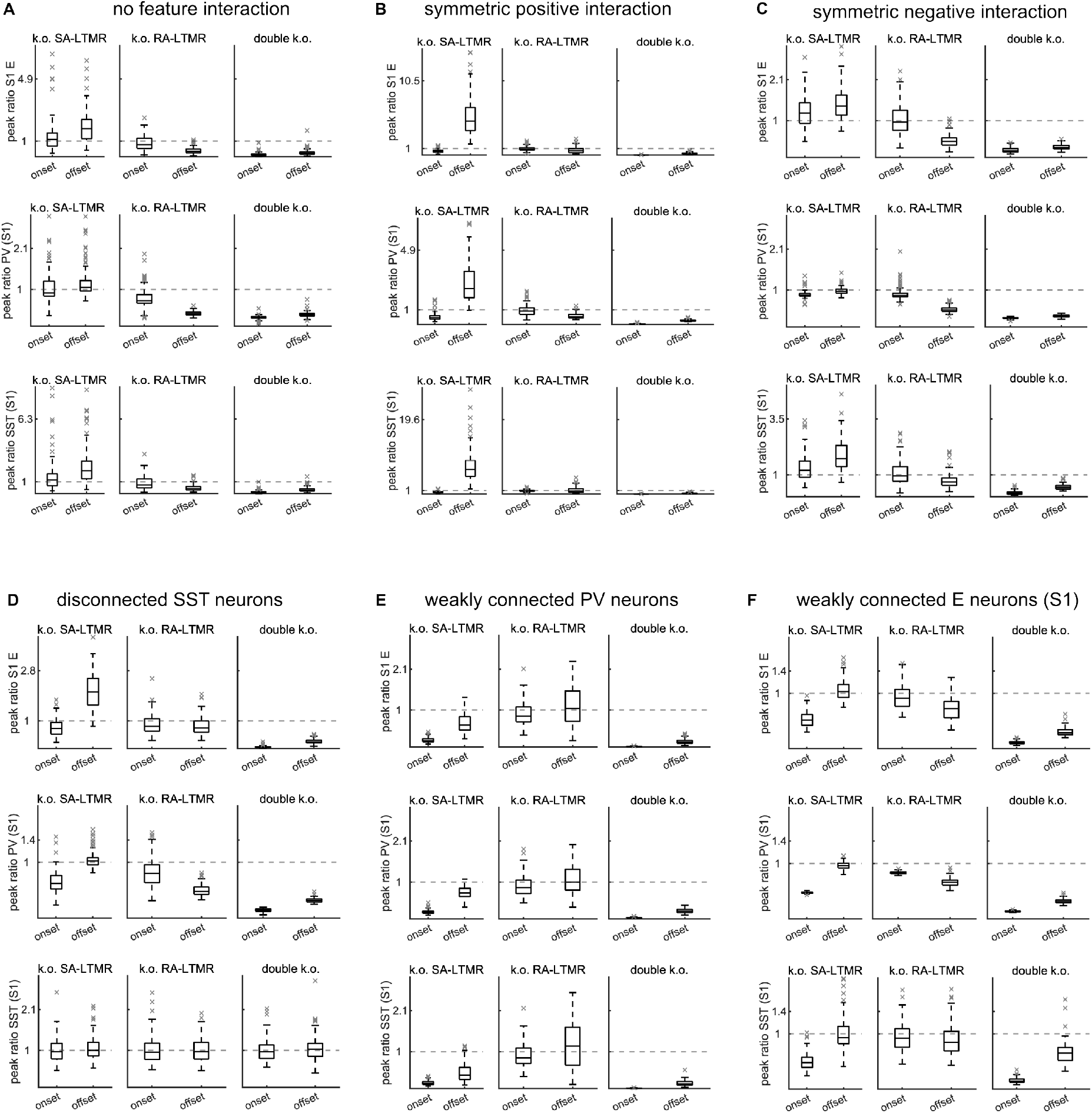
Other parameter configurations do not reproduce differential effect of receptor k.o. on firing rate in S1. **A** Same as in Fig. S5A-C, without feature interaction (*A*^(2)^ = **Id**). **B** Same as in A, with symmetric positive interaction across the fast and slow feature 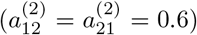. **C** Same as in A, with symmetric negative interaction 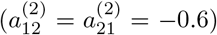. **D** Same as in A, with disconnected SST neurons. **F** Same as in A, with weakly connected PV neurons. **G** Same as in A, with weakly connectedE neurons in S1.

## References

1. Barlow, H. B. Possible principles underlying the transformation of sensory messages. Sensory Communication 1, 217–233 (1961).

2. Niven, J. E. Neuronal energy consumption: biophysics, efficiency and evolution. Current Opinion in Neurobiology 41, 129–135 (2016).

3. Olshausen, B. A. & Field, D. J. Emergence of simple-cell receptive field properties by learning a sparse code for natural images. Nature 381, 607–609 (1996).

4. Boerlin, M., Machens, C. K. & Denève, S. Predictive coding of dynamical variables in balanced spiking networks. PLOS Computational Biology 9, e1003258 (2013).

5. Bourdoukan, R., Barrett, D., Deneve, S. & Machens, C. K. Learning optimal spike-based representations. In Pereira, F., Burges, C., Bottou, L. & Weinberger, K. (eds.) Advances in Neural Information Processing Systems, vol. 25 (Curran Associates, Inc., 2012).

6. Brendel, W., Bourdoukan, R., Vertechi, P., Machens, C. K. & Denéve, S. Learning to represent signals spike by spike. PLOS Computational Biology 16, e1007692 (2020).

7. Koren, V. & Panzeri, S. Biologically plausible solutions for spiking networks with efficient coding. In Koyejo, S. et al. (eds.) Advances in Neural Information Processing Systems, vol. 35, 20607–20620 (Curran Associates, Inc., 2022).

8. Bialek, W., Rieke, F., de Ruyter van Steveninck, R. R. & Warland, D. Reading a neural code. Science 252, 1854–1857 (1991).

9. Zhu, M. & Rozell, C. J. Visual nonclassical receptive field effects emerge from sparse coding in a dynamical system. PLOS Computational Biology 9, e1003191 (2013).

10. Lochmann, T., Ernst, U. A. & Deneve, S. Perceptual inference predicts contextual modulations of sensory responses. Journal of neuroscience 32, 4179–4195 (2012).

11. Koren, V. & Denève, S. Computational account of spontaneous activity as a signature of predictive coding. PLOS Computational Biology 13, e1005355 (2017).

12. Kadmon, J., Timcheck, J. & Ganguli, S. Predictive coding in balanced neural networks with noise, chaos and delays. In Larochelle, H., Ranzato, M., Hadsell, R., Balcan, M. & Lin, H. (eds.) Advances in Neural Information Processing Systems, vol. 33, 16677–16688 (Curran Associates, Inc., 2020).

13. Golkar, S., Tesileanu, T., Bahroun, Y., Sengupta, A. & Chklovskii, D. Constrained predictive coding as a biologically plausible model of the cortical hierarchy. In Koyejo, S.et al. (eds.) Advances in Neural Information Processing Systems, vol. 35, 14155–14169 (Curran Associates, Inc., 2022).

14. Zylberberg, J., Pouget, A., Latham, P. E. & Shea-Brown, E. Robust information propagation through noisy neural circuits. PLOS Computational Biology 13, e1005497 (2017).

15. Valente, M. et al. Correlations enhance the behavioral readout of neural population activity in association cortex. Nature Neuroscience 24, 975–986 (2021).

16. Koren, V., Bondanelli, G. & Panzeri, S. Computational methods to study information processing in neural circuits. Computational and Structural Biotechnology Journal 21, 910–922 (2023).

17. Kafashan, M. et al. Scaling of sensory information in large neural populations shows signatures of information-limiting correlations. Nature Communications 12, 473 (2021).

18. Kira, S., Safaai, H., Morcos, A. S., Panzeri, S. & Harvey, C. D. A distributed and efficient population code of mixed selectivity neurons for flexible navigation decisions. Nature Communications 14, 2121 (2023).

19. Abraira, V. E. et al. The cellular and synaptic architecture of the mechanosensory dorsal horn. Cell 168, 295–310 (2017).

20. Isaacson, J. S. & Scanziani, M. How inhibition shapes cortical activity. Neuron 72, 231–243 (2011).

21. Corniani, G., Casal, M. A., Panzeri, S. & Saal, H. P. Population coding strategies in human tactile afferents. PLOS Computational Biology 18, e1010763 (2022).

22. Abraira, V. E. & Ginty, D. D. The sensory neurons of touch. Neuron 79, 618–639 (2013).

23. Handler, A. & Ginty, D. D. The mechanosensory neurons of touch and their mechanisms of activation. Nature Reviews Neuroscience 22, 521–537 (2021).

24. Neubarth, N. L. et al. Meissner corpuscles and their spatially intermingled afferents underlie gentle touch perception. Science 368, eabb2751 (2020).

25. Emanuel, A. J., Lehnert, B. P., Panzeri, S., Harvey, C. D. & Ginty, D. D. Cortical responses to touch reflect subcortical integration of ltmr signals. Nature 600, 680–685 (2021).

26. Chirila, A. M. et al. Mechanoreceptor signal convergence and transformation in the dorsal horn flexibly shape a diversity of outputs to the brain. Cell 185, 4541–4559 (2022).

27. Aronoff, R. et al. Long-range connectivity of mouse primary somatosensory barrel cortex. European Journal of Neuroscience 31, 2221–2233 (2010).

28. Pouget, A., Dayan, P. & Zemel, R. Information processing with population codes. Nature Reviews Neuroscience 1, 125–132 (2000).

29. Petersen, R. S., Panzeri, S. & Diamond, M. E. Population coding of stimulus location in rat somatosensory cortex. Neuron 32, 503–514 (2001).

30. Saxena, S. & Cunningham, J. P. Towards the neural population doctrine. Current Opinion in Neurobiology 55, 103–111 (2019).

31. Rigotti, M. et al. The importance of mixed selectivity in complex cognitive tasks. Nature 497, 585–590 (2013).

32. Fusi, S., Miller, E. K. & Rigotti, M. Why neurons mix: high dimensionality for higher cognition. Current Opinion in Neurobiology 37, 66–74 (2016).

33. Musall, S., Kaufman, M. T., Juavinett, A. L., Gluf, S. & Churchland, A. K. Single-trial neural dynamics are dominated by richly varied movements. Nature Neuroscience 22, 1677–1686 (2019).

34. Stringer, C., Pachitariu, M., Steinmetz, N., Carandini, M. & Harris, K. D. High-dimensional geometry of population responses in visual cortex. Nature 571, 361–365 (2019).

35. Stringer, C. et al. Spontaneous behaviors drive multidimensional, brainwide activity. Science 364, eaav7893 (2019).

36. Campagnola, L. et al. Local connectivity and synaptic dynamics in mouse and human neocortex. Science 375, eabj5861 (2022).

37. McCormick, D. A., Connors, B. W., Lighthall, J. W. & Prince, D. A. Comparative electrophysiology of pyramidal and sparsely spiny stellate neurons of the neocortex. Journal of Neurophysiology 54, 782–806 (1985).

38. Tremblay, R., Lee, S. & Rudy, B. Gabaergic interneurons in the neocortex: From cellular properties to circuits. Neuron 91, 260–292 (2016).

39. Wang, X.-J. Probabilistic decision making by slow reverberation in cortical circuits. Neuron 36, 955–968 (2002).

40. Meyer, H. S. et al. Inhibitory interneurons in a cortical column form hot zones of inhibition in layers 2 and 5a. Proceedings of the National Academy of Sciences 108, 16807–16812 (2011).

41. Chettih, S. N. & Harvey, C. D. Single-neuron perturbations reveal feature-specific competition in V1. Nature 567, 334–340 (2019).

42. Panzeri, S., Moroni, M., Safaai, H. & Harvey, C. D. The structures and functions of correlations in neural population codes. Nature Reviews Neuroscience 23, 551–567 (2022).

43. Runyan, C. A. et al. Response features of parvalbumin-expressing interneurons suggest precise roles for subtypes of inhibition in visual cortex. Neuron 67, 847–857 (2010).

44. Koren, V., Blanco-Malerba, S., Schwalger, T. & Panzeri, S. Structure, dynamics, coding and optimal biophysical parameters of efficient excitatory-inhibitory spiking networks. bioRxiv (2024).

45. Wilson, N. R., Runyan, C. A., Wang, F. L. & Sur, M. Division and subtraction by distinct cortical inhibitory networks in vivo. Nature 488, 343–348 (2012).

46. Kuan, A. T. et al. Synaptic wiring motifs in posterior parietal cortex support decision-making. Nature 627, 367–373 (2024).

47. Ko, H. et al. Functional specificity of local synaptic connections in neocortical networks. Nature 473, 87–91 (2011).

48. Pala, A. & Petersen, C. C. In vivo measurement of cell-type-specific synaptic connectivity and synaptic transmission in layer 2/3 mouse barrel cortex. Neuron 85, 68–75 (2015).

49. Pho, G. N., Goard, M. J., Woodson, J., Crawford, B. & Sur, M. Task-dependent representations of stimulus and choice in mouse parietal cortex. Nature Communications 9, 2596 (2018).

50. Mensi, S. et al. Parameter extraction and classification of three cortical neuron types reveals two distinct adaptation mechanisms. Journal of Neurophysiology 107, 1756–1775 (2012).

51. Niell, C. M. & Stryker, M. P. Highly selective receptive fields in mouse visual cortex. Journal of Neuroscience 28, 7520–7536 (2008).

